# Long term development of a realistic and integrated socio-ecological system

**DOI:** 10.1101/823294

**Authors:** C. Gaucherel, C. Carpentier, I.R. Geijzendorffer, F. Pommereau

## Abstract

We developed a discrete and qualitative model of integrated socio-ecosystems, with the help of formal Petri nets. We illustrated such Petri nets in the case study of temporary marshes in the Mediterranean part of France, the Camargue delta, by integrating biotic, abiotic and human-related components along with their processes into the same interaction network. The model demonstrated that when marshes are exposed to extensive grazing the presence of marsh heritage species is facilitated by opening up the vegetation through various trajectories. This supports the commonly used management practice of extensive grazing to conserve certain protected habitats. With this Possibilistic approach, we identified all potential ecosystem trajectories and provided their differential (non-systematic) impacts on heritage species richness (number). Hence, we rigorously demonstrate with this new type of model that grazing benefits marsh species which are faced with competition from common grassland species. The detailed analysis of the explicit state space and trajectories allows exploring simultaneously the identification of a range of recommendations for management strategies.

## Introduction

Historically, ecosystems have been predominantly studied through a focus on species communities embedded into the environment, thereby reducing the abiotic part to a simplified resource input or as an external source of perturbations (e.g. Kéfi et al. 2016; Thébault and Fontaine 2010). Yet, abiotic components are an intrinsic part of the biophysical and socio-economical entity that defines an ecosystem, thus playing a key role in its dynamics through various feedbacks (Frontier et al. 2008; Gaucherel 2019). Nowadays most ecologists are convinced that an integration of all abiotic, biotic and even human components is required to improve our understanding of ecosystem functioning.

In the context of impacts of global changes, biophysical interactions and the influence people have on ecosystems, are transforming the ecosystem structure, sometimes drastically so (Cincotta et al. 2000; Cumming et al. 2014; Ostrom 2009). Unfortunately, despite a common assumption that integration of abiotic, biotic and human components is required to understand how ecosystem structure may be affected, attempts in this line are considerably outnumbered by studies dedicated to simplified systems or to single elements (but see Geijzendorffer et al. 2017; Titeux et al. 2017). In addition, the integration of components of multiple dimensions also enlarges the window of potential ecosystem transitions to an extent that is difficult to manage in a computational and comprehensible manner. In this paper, we propose a conceptual and methodological framework to develop a generic model of ecosystem *functioning* and *development* (i.e. structural changes over the long term, Gaucherel et al. 2017).

Modeled biotic interactions have been defined as trophic or non-trophic ones (Thébault and Fontaine 2010), parasitic or pollinic ones (Campbell et al. 2011), competition or facilitation (Lefever and Lejeune 1997), or, more rarely, a combination of several kinds simultaneously. Recently, multiplexes have been proposed to represent several types of interactions between the same species nodes within the same ecological network (Brose et al. 2006; Kéfi et al. 2016), thereby strongly simplifying the overall ecosystem. Multiplexes are models that intend to grasp a subset of processes that play a dominant role in the overall system dynamics (fates) and, in particular, to study the effects of interactions (i.e. processes between two or more system components) on the system stability.

When not limiting system models to species interactions only, they could also be used to integrate abiotic, biotic and human components and related processes to study their combined impact on socio-ecosystem components and structures (i.e. the shape of the interaction network). The model can then be used to study trajectories (i.e. pathways combining successive atomic transitions) of these systems under different scenarios. Although this may pose methodological challenges, it seems feasible to develop a more holistic interaction network (i.e. made up of a large number of biotic, abiotic and human components and their related processes). Knowledge on how to develop integrated networks is already available with parameterisation of the model being a known hurdle (Frontier et al. 2008; Gaucherel et al. 2019; Ricklefs and Miller 2000). To handle potentially huge and complex interaction networks, new kinds of models and/or methods of simplifications are required. Here, we demonstrate two different ways to simplify the study of ecosystem trajectories: first, we model qualitative interaction networks, as already explored in biology (Reisig 2013); secondly, we focus only on the structural changes of this network (Gaucherel and Pommereau 2019; Gaucherel et al. 2017). In other words, we model network (topological) changes instead of dynamics (fluxes) within a frozen network.

The objective of this study was to explore the diversity of trajectories of a socio-ecosystem structure that becomes possible when integrating abiotic, biotic and human components. Depending on the way an interaction network has been defined, structural changes best represent long term dynamics like the arrival of invasive species or pollutants, species extinctions or human migrations. Here, we illustrate the “ecosystem development” concept for a vulnerable ecosystem within the European habitat Directive; the Mediterranean temporary marshes, located in the Camargue, South of France (Beltrame et al. 2013; Grillas and Roché 1997) and their conservation poses several challenges (Medail et al. 1998; Rhazi et al. 2006). Temporary marshes are fragile ecosystems often subjected to grazing as part of the management (Beltrame et al. 2013; Duncan 1992). The general current understanding being that grazing reduces the cover of dominant species, which are often common species, thereby creating space for new or rare species (Chambers and Prepas 1990; Gough and Grace 1998). However, this understood impact of grazing on vegetation greatly depends on the intensity of the grazing (Noy-Meir et al. 1989; Sternberg et al. 2000) and negative conservation impacts from grazing regimes are not uncommon (Bouahim et al. 2010). An increased understanding of the impact of grazing on these fragile ecosystems would allow to adjust the conservation management plans (Beltrame et al. 2013; Duncan 1992).

To develop an integrated network model of these marshes, we borrowed a formalism (a mathematical framework) from theoretical computer sciences which was conceived to handle large network changes: a qualitative Petri net. Petri nets belong to the wide category of discrete modelling formalisms (Giavitto and Michel 2003; Pommereau 2010). Such models have proven powerful when applied to biological networks but, to our knowledge, are almost absent in ecology (but see Baldan et al. 2015; Gaucherel and Pommereau 2019; Gaucherel et al. 2017). We demonstrate the ability of Petri nets to integrate abiotic, biotic and human components of a system and to handle its sharp changes over the long term. For this purpose, we defined two contrasting scenarios of temporary marsh systems: marsh dynamics without and with extensive grazing. In a reductionist view, we listed all the extensive grazing impacts (e.g. trampling, browsing or fertilizing effects) possibly observed on various marsh vegetation types, thus avoiding more intensive actions. We then tested the hypothesis that, in a holistic (integrated) view of the system, grazed marshes would favour the presence of typical marsh biodiversity (heritage species) in comparison to non-grazed marshes, in good agreement with the intermediate disturbance theory (Fox 2013; Wilkinson 1999). Additionally, we demonstrated that our qualitative model is able to identify a diverse and comprehensive range of trajectories (also called the “development”, i.e. all future states and structural changes) of the integrated socio-ecosystem which allow for the identification for a rich set of management recommendations. We finally discussed the properties of this innovative approach.

## Materials and Methods

### Temporary marshes in Camargue

Temporary Mediterranean marshes are defined as small shallow depressions (<10 ha and less than 2m depth) fed by precipitation in autumn or spring (Fig. 1a). These ecosystems are characterized by a natural alternation of flooding and drying out phases providing a niche for many aquatic, amphibian and terrestrial species (Grillas et al. 2004). Temporary marsh allows the development of an adapted flora rich in aquatic (*e.g. Zannichellia sp.)* and amphibious (*e.g. Damasonium polyspermum*) species, thus totally dependent on water dynamics (Grillas and Roché 1997). Similarly, the temporary marsh fauna usually show a two-phase life cycle (Grillas et al. 2004), such as for remarkable amphibian species (*e.g. Pelobates cultipes*) and dragonflies (*e.g. Lestes macrostigma*), while ichthyofauna is absent (Grillas and Roché 1997). Moreover, the abundance of plant and animal resources in these marshes make them a feeding habitat for many birds (*e.g. Anas crecca*), as well as for wild mammals (*e.g. Sus scrofa*) and domestic ones (e.g. cattle).

**Figure 1.**
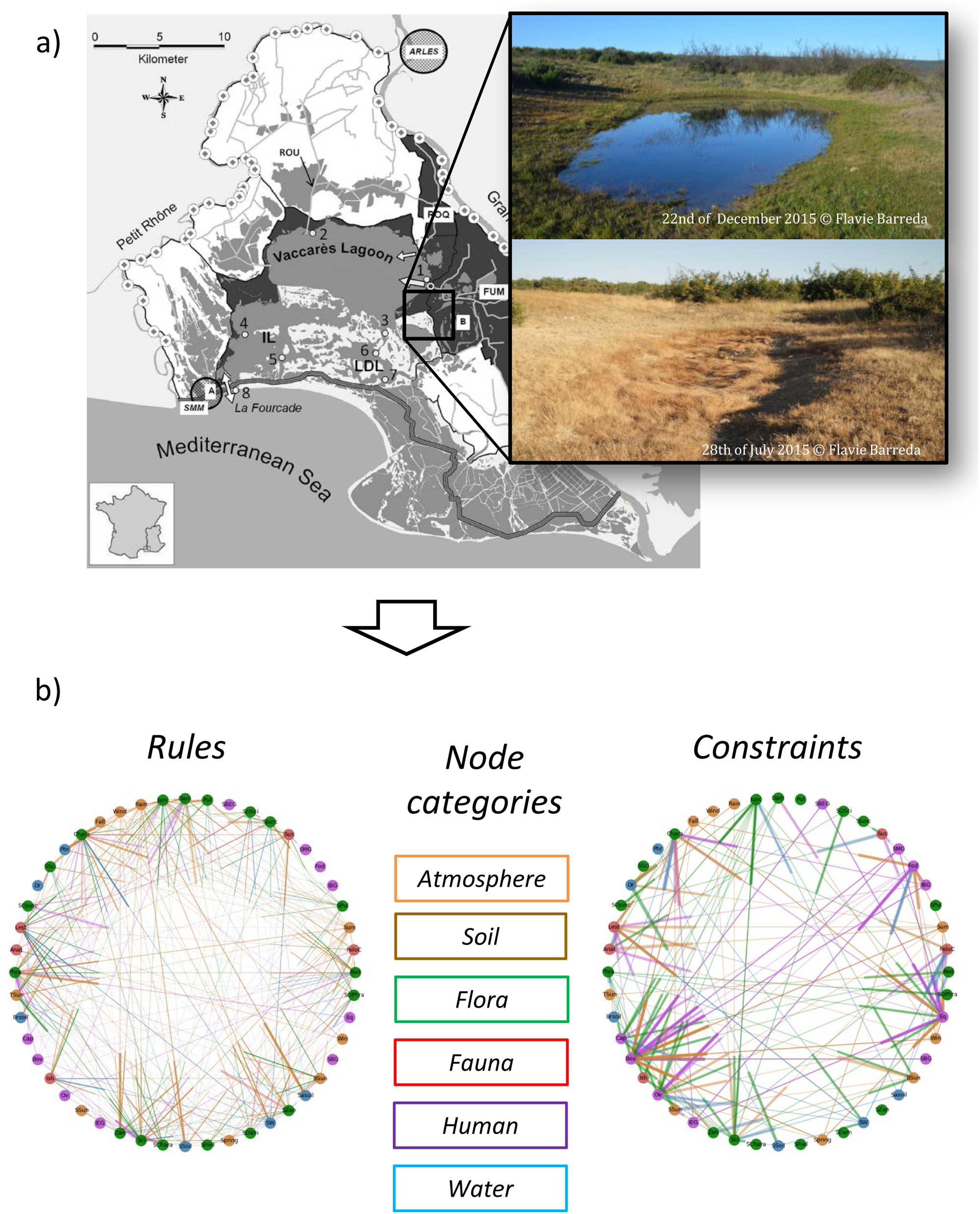
Diagram of the Camargue delta (a), with photographs of a temporary marsh, and the ecosystem graphs (b) defining the temporary marsh model. The Camargue is located between the Mediterranean sea and the city of Arles (a), south of France, and hosts numerous temporary marshes (a, insert) alternatively drying and filling up during the year. We modeled such marshes with 46 components (nodes, b) connected through 105 interactions *stricto sensu* (b, left) and 57 interaction constraints (b, right) standing for ecosystem processes (Appendix 4, Tables S5 and S6).

Domestic cattle (*e.g.* equines and bovines) graze marshes during their terrestrial phase and/or their drying out phase. The act of grazing entails various disturbances such as trampling, browsing or fertilizing inputs (Mesléard and Perennou 1996). In the Tour du Valat domain, in the Camargue delta, temporary (non-brackish) marshes account for 250 ha. Within the model, we do not consider any other major disturbance than the extensive grazing by domestic animals, excluding thereby the potential impact of invasive species and assuming no connection with permanent water bodies (Duncan 1992; Grillas and Roché 1997; Mesléard and Perennou 1996). Here, we developed a spatially implicit model (although it is possible to take into account for spatial units in the ecosystemic graph, Gaucherel et al. 2012), every processes occurring at the same place in the temporary marsh. Also, we considered relatively short term processes, namely, intra-annual changes cumulated over less than five-year duration.

### An integrated ecosystem Petri net

To test our central hypothesis, we used the species community as a proxy of ecosystem structure changes. To analyse the impacts of grazing on a subset of species uniquely linked to the temporary marshes in Camargue, we successively modelled: i) the hydrology of a temporary pool, ii) requirements of domestic animals, and iii) characteristics of species of the temporary marsh ecosystems with conservation value. We represent the ecosystem by its interaction network and with an *ecosystem graph* made of material components, the nodes, connected through their related (immaterial) processes, the edges. For each set of processes, we identified the components of interest based on literature and expert knowledge (Table 1), namely: surface water for temporary ponds, bovines and equines for domestic mammals and species of conservation value listed by the Tour du Valat (Grillas et al. 2004; Mesléard and Perennou 1996). The definition and integration of these components and their processes were populated using literature and interviews with experts at the Tour du Valat research institute (Appendix 4, Tables S5 and S6, with references therein).

**Table 1.**
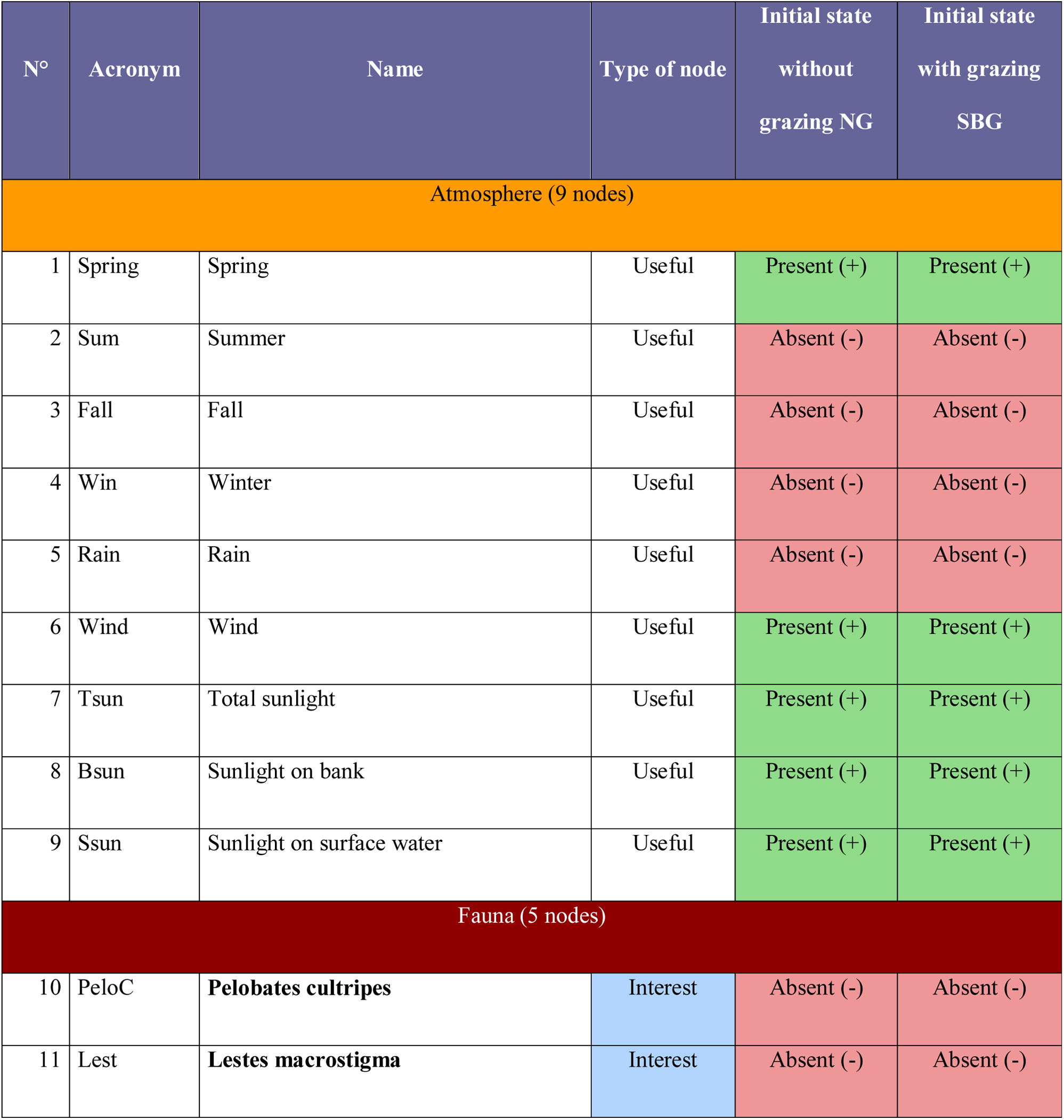
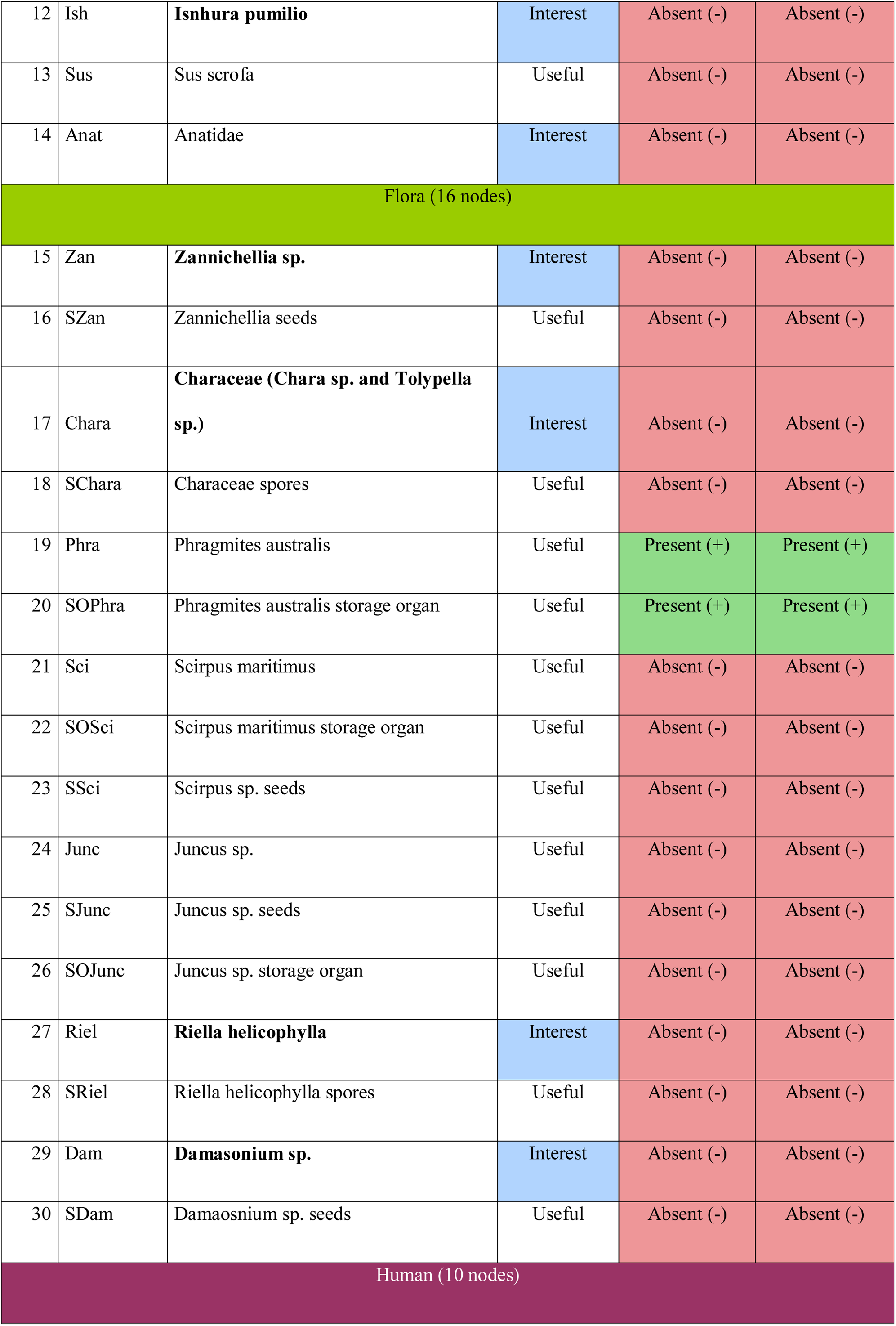
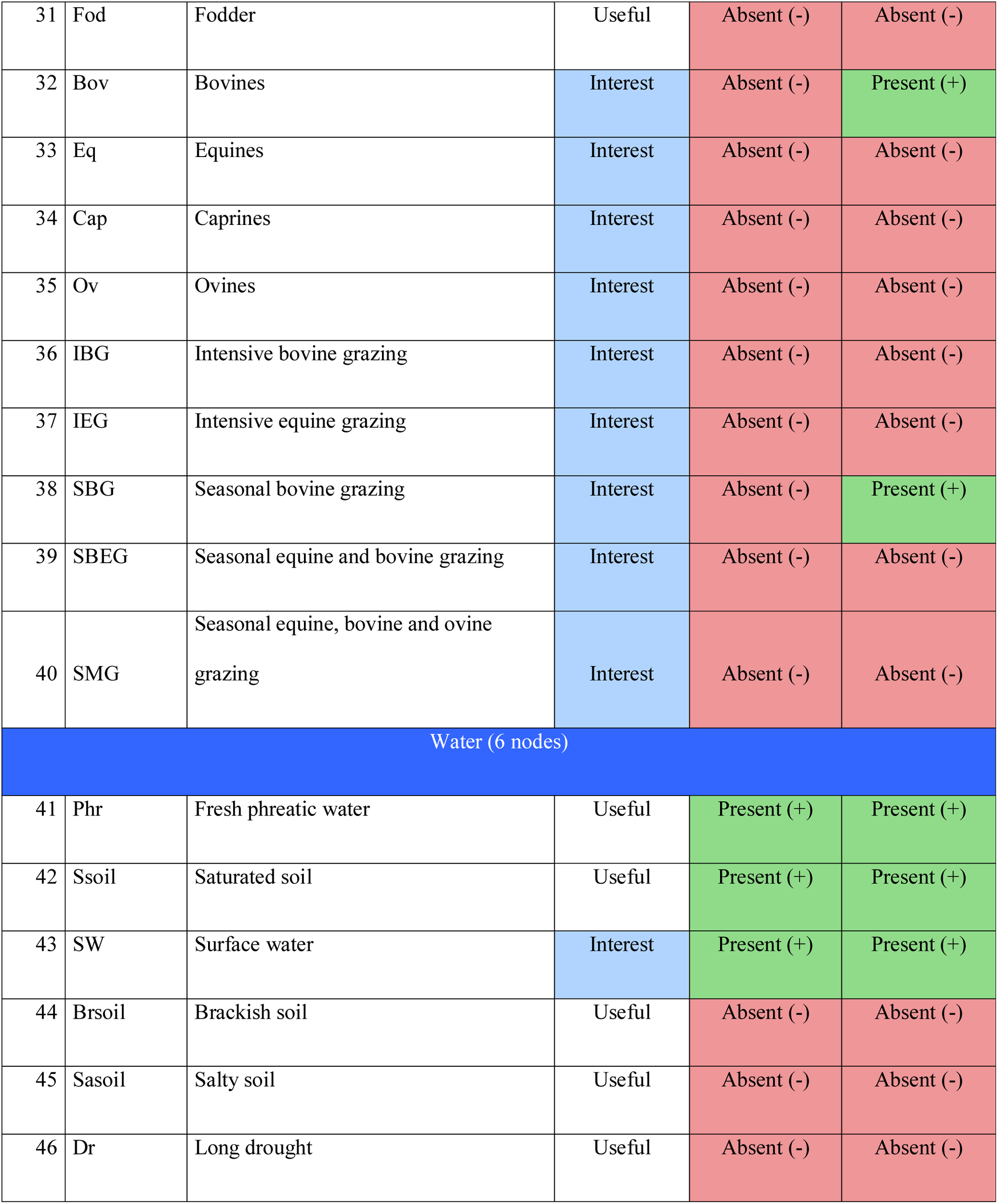
Model components and node categories, names, abbreviations, and descriptions of the temporary marsh ecosystem graph. Whether these ecological components are useful for the ecosystem functioning or of interest for the addressed question is displayed (species with conservation value are in bold), as well as the initial conditions of both NG and SBG scenarios (last columns).

This resulted in a qualitative model consisting of 46 components with 105 interactions *stricto sensu* and 57 interaction constraints, standing for mandatory processes (Fig. 1b and Appendix 3). To gain objectivity, all modelled components should be connected to other components (i.e. at least one process impacts it, and at least one process depends on it). We call such connected nodes *characterized* components (Gaucherel and Pommereau 2019). In our temporary marsh model, the best described components are those associated with the central question addressed (*i.e.* concerning species grazed by domestic animals and related species of temporary marshes (Appendix 4, Tables S5 and S6)). Finally, the remaining semi-characterized components were fixed if they had an impact on the defined ecosystem but were not influenced by it (e.g. components relative to the atmosphere and to grazing). The resulting model handles Boolean components (i.e. being functionally present or absent) and qualitative processes (being executed or not, not to be confused with Boolean functions), to ultimately explore all possible ecosystem developments in the future from an initial ecosystem composition called the *initial state* (Gaucherel and Pommereau 2019; Gaucherel et al. 2017).

### Petri Nets and a simplistic Predator-prey model

We illustrate here the functioning of the model using a simplistic predator-prey system (Fig. 2). Additional details on the principle and uses of Petri nets can be found in literature (Gaucherel and Pommereau 2019; Pommereau 2010; Reisig 2013) and in Appendix 2. The ecosystem Petri net itself is developed in three successive steps: a) an intuitive graph (i.e. a set of components and their related processes) is built to represent the studied ecosystem based on the leading question; b) we then transform this ecosystem graph into a formal model based on Boolean components and evolution rules; the components have an initial value that define the initial state; c) applying these rules on the initial state allows computing new states, causally linked, on which the processes may be repeated until no new state can be computed; this progressively yields the state space of the system; d) to avoid building the state space by ourselves, we translate the rules into a Petri net with SNAKES, a software tool (Pommereau 2015); e) the state space of the Petri net (also called its marking graph) is always strictly equivalent to the state space we would have obtained at step (c). It is worth noting that steps (d) and (e) are usually hidden to the users who directly get the state space (c) obtained from the marking graph (e).

**Figure 2.**
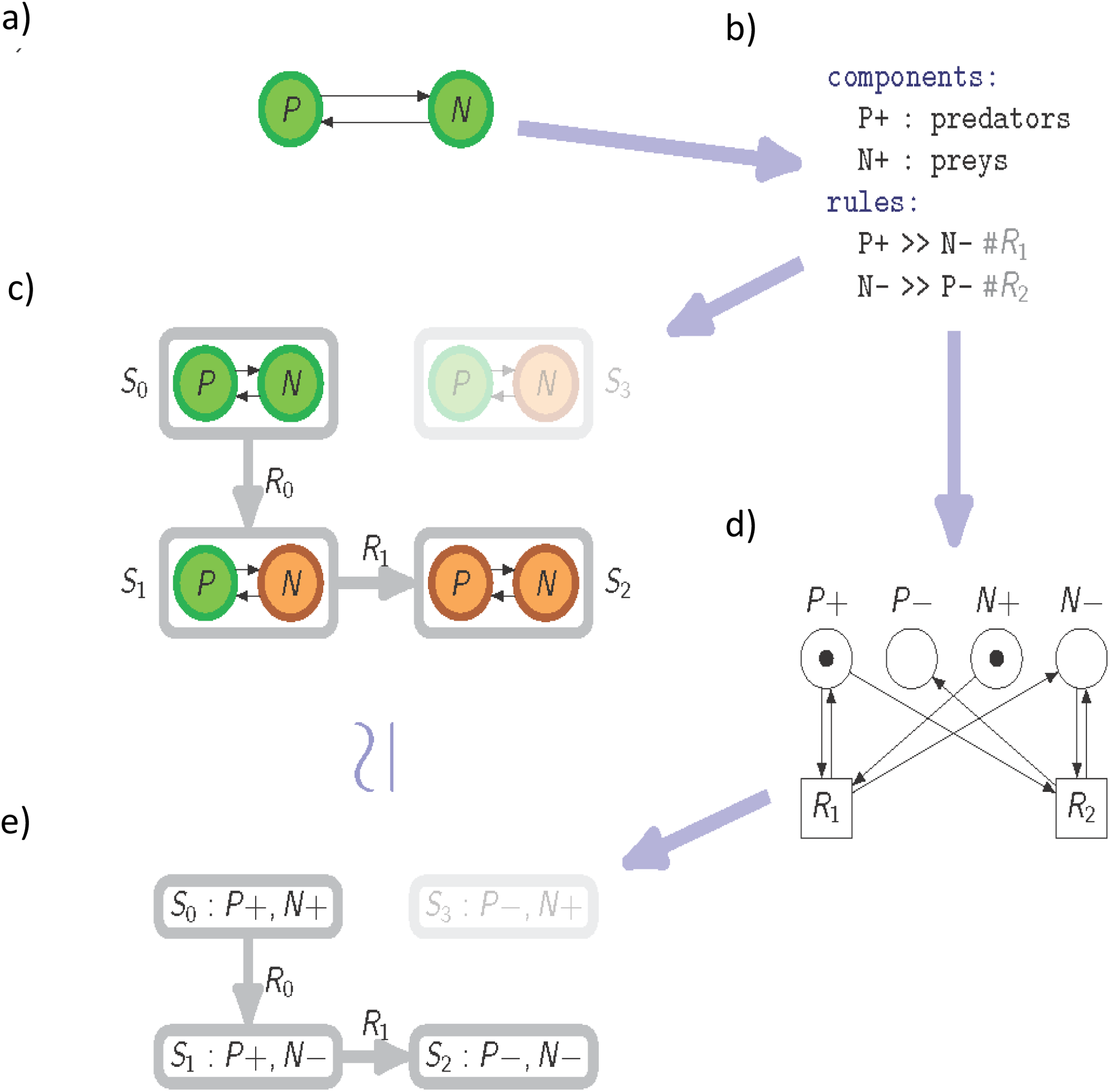
Illustration of a simplistic predator-prey system (a), with its associated Petri net (b), its qualitative dynamics (c), and marking graph (d). The system is made of two components, the prey (N) and predator (P) populations, and two interactions connecting them (rules R1 and R2), as seen on the automaton (a). The corresponding Petri net is made of four places (P+, P-, N+, N-) and two transitions R1 and R2, where unlabeled arcs have weight 1 (b). Starting with the presence of both populations, it is possible to list all system states encountered (c), and to connect them with the rules (absent nodes and inactivated rules are displayed in grey). The net is depicted in the initial state (c), and the successive states may be deduced from the token circulation seen in the dynamics (b). The marking graph of the Petri net (d) is depicted with each state number (S_0_, S_1_, S_2_) referring to the dynamics described above (b). Notice that a specific state of the system (S_3_) may not be reached from this initial condition and with these rules (d).

Any ecosystem can be represented as a multi-digraph (i.e. a directed graph with parallel edges). In this graph, every material component of the ecosystem (e.g. abiotic: precipitation; biotic: species; human: domestic cattle) is represented by a *node*, with two Boolean states: “present” (the component is functionally present in the system, also denoted “+” or ON) and “absent” (functionally absent of the system or “-” or OFF). In the simplistic example predator-prey system only two nodes are defined: the prey and the predator populations. Any *state of the system* is defined by the set of “+” and “-” nodes (Fig. 2b). The maximal number of possible system states is 2^#nodes^ and grows exponentially with the number of nodes. The *rules* correspond to any physicochemical, bio-ecological and/or socio-economic processes (e.g. if there are no preys there are no more predators), and thus represent all possible interactions between nodes composing the ecosystem studied. In the predator-prey system, two rules only are defined: R1, the predation itself: the predator eats the prey, and R2, the mortality: without prey, the predator dies (Fig. 2b and 2c). In the Petri net language, nodes are called places and rules are called transitions, both being connected through (oriented) arcs (Fig. 2d).

A separate rule describes the *condition* and a *realization* parts as: “transition’s name: condition ≫ realization” (see Appendix 2 for formal definitions). For a rule to be applied the state of the node must satisfy its application condition; the rule is then said to be “*enabled*”. If so, the application of the rule modifies the state of the nodes as stipulated in its realization part; the rule is *fired*. The firing of a rule, including testing its condition, is always atomic (i.e. no other rule can interfere during the firing). In the predator-prey system, the rules are written as *R1: P*+, *N*+ ≫ *N-* and *R2: N-, P*+ ≫*P-* (Fig 2b). Since the rules modify node states, they change the overall system state (i.e. the state of the system aggregates all node states). Therefore, the entire system shifts from one state to another one through the discrete successive application of rules (Fig. 2c). The repeated execution of the rules progressively produces the *state space*, which provides the set of all system states reachable by the rules defined (Fig. c); this can be equivalently (and more conveniently) obtained through the Petri net and its marking graph. As a corollary, the system states are connected to each other by some of these rules in the state space too. The size of this state space is usually much smaller than the number of possible system states, because the computation starts from a specific initial condition and because rules have specific application conditions. We develop some tools to automatically compartmentalize large state spaces into *merged* (simplified) state spaces (Appendix 2).

Firing a rule independently to some others often leads to unrealistic trajectories (e.g. removing water without removing fishes in it). Therefore, we defined particular rules, called *constraints*, preventing the model from reaching such unrealistic (socio-ecological) trajectories. Constraints have a condition and a realisation part, just as rules *stricto sensu* do, and model inevitable (mandatory) events given the system state. The sole difference between rules and constraints is that constraints have priority on rules *stricto sensu*. In the predator-prey system, the system state S_1_ = (N-, P+) may be considered as unrealistic; so, the rule R2 may be transformed into a constraint (*C1: N-, P*+ ≫*P-*). From a given state, the model first computes all trajectories opened up by the defined constraints and then only, when all the system states obtained are realistic (*i.e.* there is no longer any enabled constraint), the enabled rules are fired (Fig. 2c). In brief, the discrete model proposed here is qualitative, mechanistic (processes are explicit), deterministic (no stochasticity but with several outcomes) and asynchronous (all rules are applied as soon as possible, no rule conflict) (Gaucherel and Pommereau 2019; Reisig 2013).

### Methodology and scenarios

The impact of grazing on the marsh species is studied by comparing two contrasting scenarios, namely no grazing (noted NG for non-grazing) and seasonal extensive grazing by bovines (SBG for seasonal bovine grazing). The latter scenario corresponds to extensive bovine grazing during spring and summer, and to the absence of bovine in winter and fall (seasons during which bovines graze more productive lands, Duncan 1992; Grillas and Roché 1997). For an easier comparison, each scenario is simulated on the basis of the same sets of nodes and rules; only their initial states differ. This allows some rules to be enabled in one scenario but not in the other one, thus corresponding to contrasted *development* for the same ecosystem, i.e. contrasted NG and SBG sequences of states. As a consequence, each scenario leads to a specific state space. Since both scenarios are based on the same nodes and rules, a subset of system states will be in common (except for the node modified in the initial state). We will first plot the histograms and basic statistics of presence/absence counts of the marsh species (ranging from zero to a maximum of seven species with conservation value) in each state space to quantitatively visualise the similarity and differences between both scenarios.

Both (NG and SBG) initial states consider a temporary marsh to contain water in spring, with a *Phragmites australis* population. The initial state of the SBG scenario has two more present nodes: “Bovine” and “Seasonal bovine grazing”, fixed and semi-characterized nodes. This initial state allows four constraints to be enabled at each season shift (Appendix 4, Tables S5 and S6). As each state space is explicitly computed (i.e. all states are known and described), we may compute the present and absent nodes in each state and look for states satisfying specific criteria (Appendix 2). In addition, we developed specific tool to *merge* both (NG and SBG) *full* state spaces (Appendix 2) and to automatically compute the strongly connected components (SCCs) of state spaces, which are structural stabilities gathering interconnected ecosystem states. All states of the final merged state space are analyzed in terms of vegetation types and average number of marsh species with conservation value. In such analyses, we will plot both NG and SBG merged spaces colored by averaged species numbers and by (downward) location relatively to the initial state. Finally, we will perform two principal component analyses (PCAs) on the two merged state spaces to analyze details in the dominant (merged) trajectories reached by the ecosystem.

In the modelling approach presented here, every *material* component involved into the long term changes of the ecosystem is represented as a node, while every (immaterial) process is represented as an oriented edge between two or more nodes. Such an informational conception of ecosystems is based on a theoretical view still under test (Gaucherel et al. 2019). The Petri net engine allows to mathematically formalize the topological changes of the integrated interaction network (Gaucherel et al. 2017). By this way, every ecological process (e.g. predation, parasitism, pollination or competition) has a formal and qualitative definition whose consequences in the network changes can be rigorously assessed, and merged with others (Fig. 2). In a second step, this approach allows identifying all possible trajectories of the studied systems (Gaucherel and Pommereau 2019), by handling the node and edge changes (the topology) of the interaction network. As the state space of the system is equivalent to the defined Petri net (Gaucherel and Pommereau 2019), it becomes possible to explore all reachable trajectories of the ecosystem, and then to identify sustainable or potentially unwanted trajectories. The model provides results that resemble those of differential equations, while equations focus on singular, statistically most probable trajectories only (Dambacher et al. 2003; Thébault and Fontaine 2010). In addition, our qualitative model strongly differ to others by the fact that a set of rules differs from functions, not providing a unique value output, and possibly grasping conflicting and even opposite responses of variables (nodes).

## Results

### State space and general analyses

The initial states NG and SBG lead to two different states spaces showing 1.6 and 1.54 10^6^ states, respectively (Fig. 3a). Both state spaces have 1.19 10^6^ states in common, hereafter called the reference set. By removing this reference set, we identified exclusive states for the scenario without grazing (NG → 4.04 10^5^ states) and with seasonal bovine grazing (SBG → 3.53 10^5^ states). The NG set is much more aggregated (58.59 % of the state successors of any NG state are also in this set) than the SBG set (6 % only), thus showing an exclusive dynamics. Moreover, the SBG set is associated with a significant increase in number of species with conservation value (p-val < 2^−16^, Fig. 3a).

**Figure 3.**
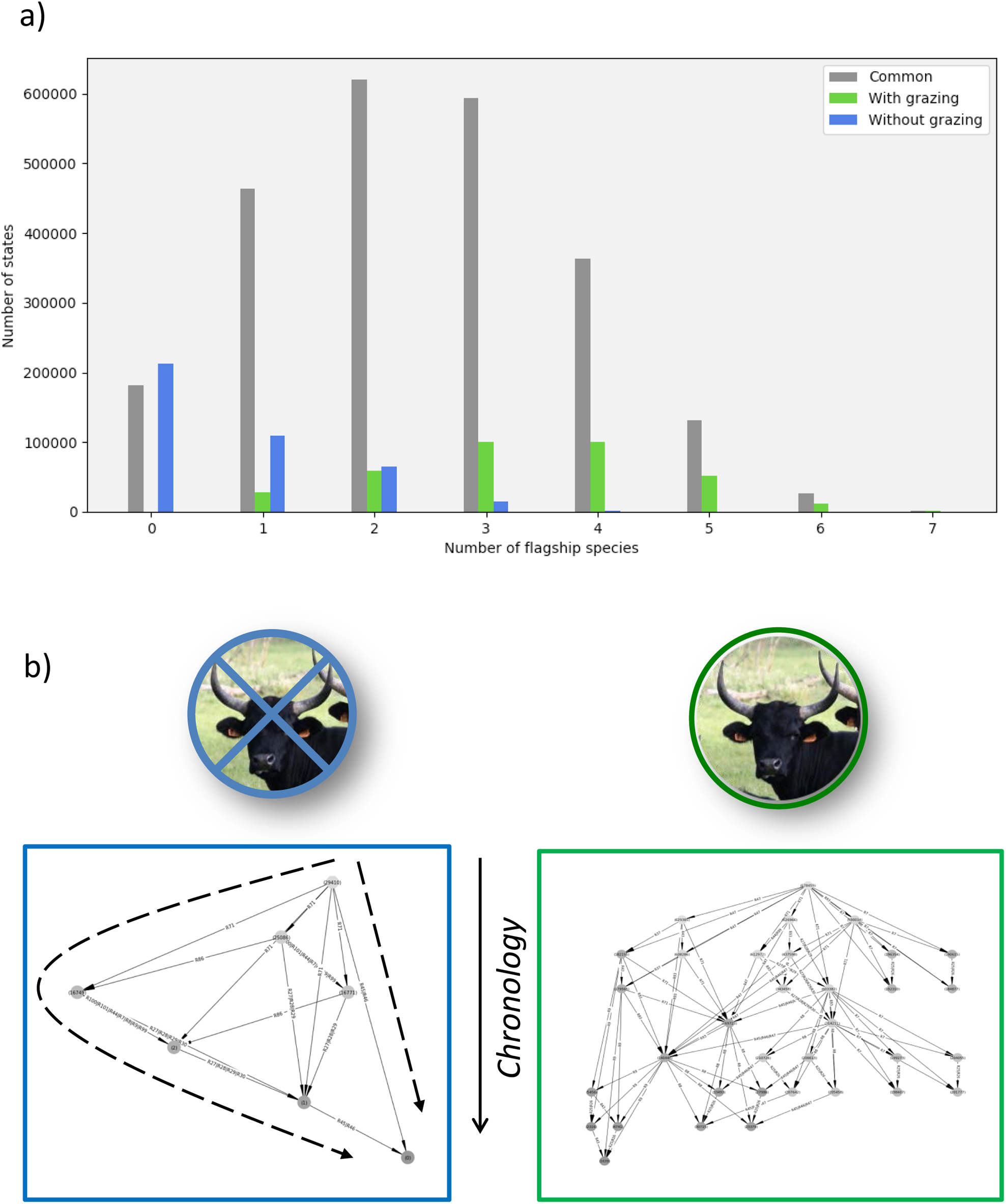
Histogram of the ecosystem states (a) when run with (green) and without (blue) grazing, and their common states (grey), along to the explicit merged state spaces (b). The histogram shows the counts of states containing heritage species (a, ranging from 0 to 7, Appendix 3, Table S2) for reference (in grey), NG (non-grazing, in blue) and SBG (seasonal bovine grazing, in green) scenarios over the full states spaces (Appendix 3, Table S4). The NG (b, left) and SBG (right) merged state spaces are displayed with colors representing the average number of heritage species in each strongly connected component (each disk gathers several highly connected states). The ecosystem trajectories should be read from top to bottom and following the arrows connecting every component (disks).

The NG set suggests a marsh undergoing a strong drying out due to the lack of bovine-induced processes: the states of this set encompass less surface water (SW, Table 1), a saturated soil (Ssoil), more brackish and saline soil (Brsoil, Sasoil) and long drought (Dr). Indeed, all these nodes appeared significantly higher in the NG set than in the reference set (p-val < 2^−16^). The SBG set is characterized by a high frequency of spring states, but this frequency does not impact the number of species with conservation value. In addition, the SBG scenario shows a decrease of the helophytes (*Scirpus maritimus, Phragmites australis* and *Juncus sp*.) and an increase of the light on the site, in opposition to the NG scenario. Indeed, the SBG set contains states with significantly higher numbers of helophytic species than in the reference set (p-val <2^−16^). The merged state spaces of NG and SBG scenarios (Fig. 3b) highlight the dominant trajectories of the ecosystem. As the model is causal, it is possible to identify trajectories with averaged compositions of strongly connected components by following downward the ecosystem chronology (Fig. 3b, dashed arrows).

### Ecosystem development and scenarios

The merged state spaces of NG and SBG scenarios highlighted the dominant trajectories of the ecosystem in terms of average species numbers in strongly connected components (colors in Fig. 4a and 5a). So, in terms of conservation needs, red instead of blue sets of states (SCCs) should preferably (and may) be avoided. The variance distribution of the state space, using a principal component analysis (PCA) performed on SCC profiles (node compositions), does not provide a simple biological interpretation for the number of species with conservation value (Fig. 4b): no univocal correlation exists between species number and marsh properties. Indeed, the NG scenario produced a merge state space made up of seven SCCs, gradually and irreversibly shifting toward a final and huge SCC with herbivory (number 0, Fig. 4a). The NG variance distribution of the PCA decreased sharply with 48.8 % (resp. 22.2 %) of the total variance obtained from the first (resp. two) main PCs. The PC#1 is closely related to climate and seasonal variations, with spring on negative values (left hand side, with wet system states) and winter, autumn (all dryer states) on positive side (Fig. 4b, right side). The PC#2 is mainly related to the marsh vegetation and helophyte types, between rushes on negative values and scirpae (*scirpus maritimus*) on positive values of this axis, whatever the seasons. When locating the SCCs in the PC1/PC2 plane (Fig. 4b), we follow the dominant trajectory of the ecosystem (downward in Fig. 4a), from the relatively dry and scirpae-type vegetation to relatively wet and richer system states, and reversely then (Fig. 4b, left side, dashed arrows). Such state spaces precisely illustrate a socio-ecosystem duality, with some PCA axes dedicated to biotic components and processes, some others to abiotic ones.

**Figure 4.**
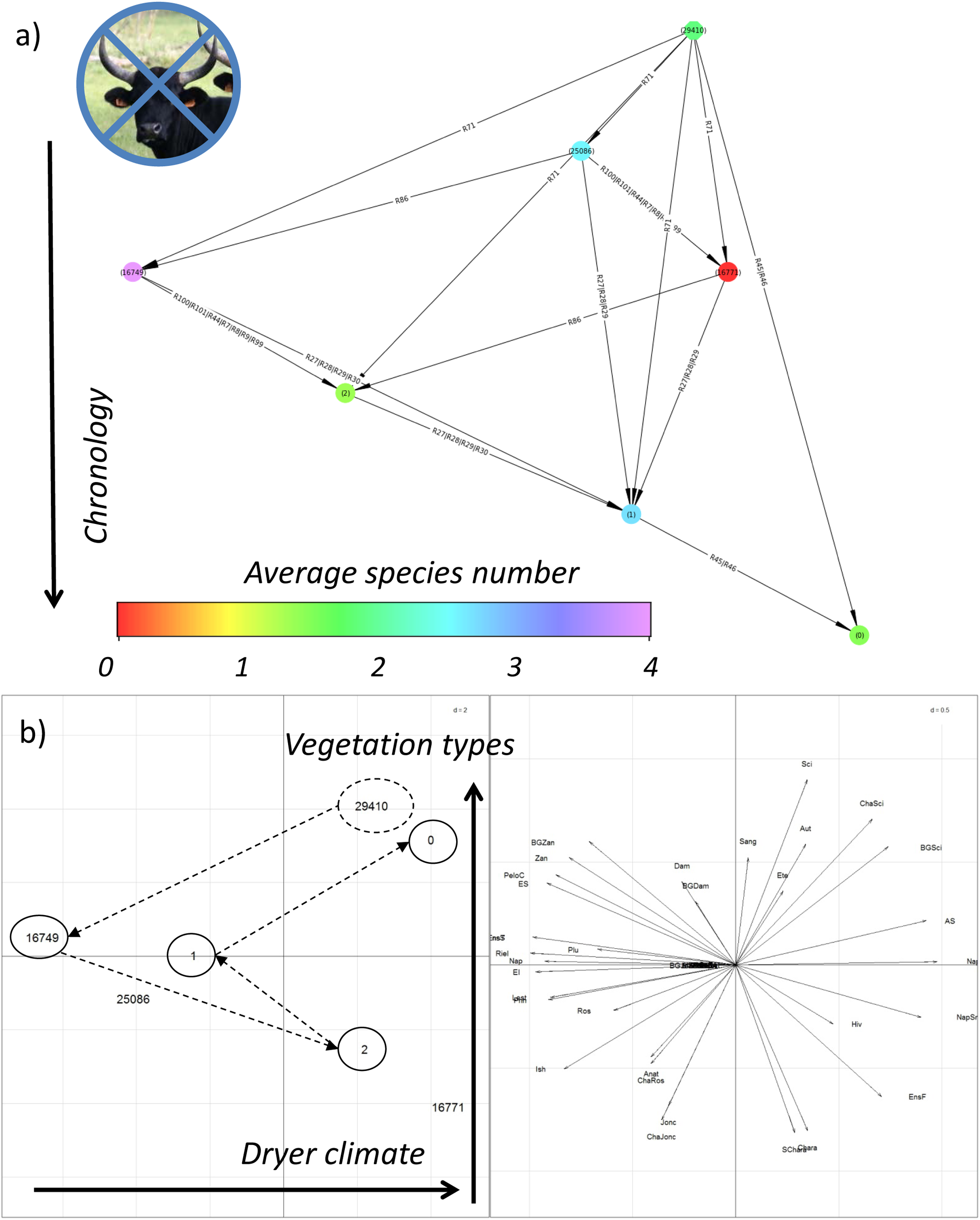
The merged state space of the NG (non-grazing) scenario (a), and its corresponding statistical analysis (PCA, b). The merged state space should be read from top to bottom, through the various trajectories connecting every strongly connected component (a, disks) with various rules (on arrows). The components are gathered (a, ellipses) according to their locations in the PC1/PC2 plan of the PCA (b, left), and each component is colored according to its location on PC1. The statistical analysis (b, right) is easily interpreted according to the projections of the ecosystem nodes involved in each component, and allows reconstructing the ecosystem trajectories (a, ellipses and dashed arrows) in terms of ecosystem composition (b, dashed arrows).

The SBG scenario produced a much more complex merge state space made up of 34 SCCs, gradually and irreversibly shifting toward four final SCC groups (#1639 group A, 30715, 23379, 207642 and 205458 group B, 198407 and 201777 group C, and 192310 and 184877 group D, Fig. 5a, ellipses). The SBG variance distribution of the PCA decreased sharply with 41.5 % (resp. 17.8 %) of the total variance obtained from the first (resp. two) main PCs (Fig. 5b). The PC#1 is closely related to climate and seasonal variations and opposite to the NG PC#1 (i.e. wet and higher species numbers). The PC#2 is mainly related to the marsh vegetation types, with reeds C on positive values, and scirpae group A on negative ones). Most ecosystem trajectories under NG are projected in the upper part of the SBG scenario merged space (not shown), thus confirming that bovines are later on reducing the marsh salinity, opening up the lower strata of vegetation and thus providing longer trajectories (i.e. with higher discrete steps) with more monospecific vegetation types (the groups). When locating these SCCs in the PC1/PC2 plane (Fig. 5b, left side), the ecosystem basically shifts from a rich and wet (spring) state on average to a relatively dry ecosystem with various vegetation type. Yet, going back to the corresponding state space colored by the number of species with conservation value (Fig. 5a), final SCC groups are not systematically the poorest states, or the richest ones. So, optimal trajectories may be selected on this basis, in particular focusing on some decisive SCCs (SCCs number 590014, 314211 and 140448, Fig. 5b arrows) from which the ecosystem irreversibly shifts toward some specific and less diversified groups (ellipses). Indeed, the model demonstrates that the system may not recover previous states after reaching these so-called terminal SCCs (Fig. 5a).

**Figure 5.**
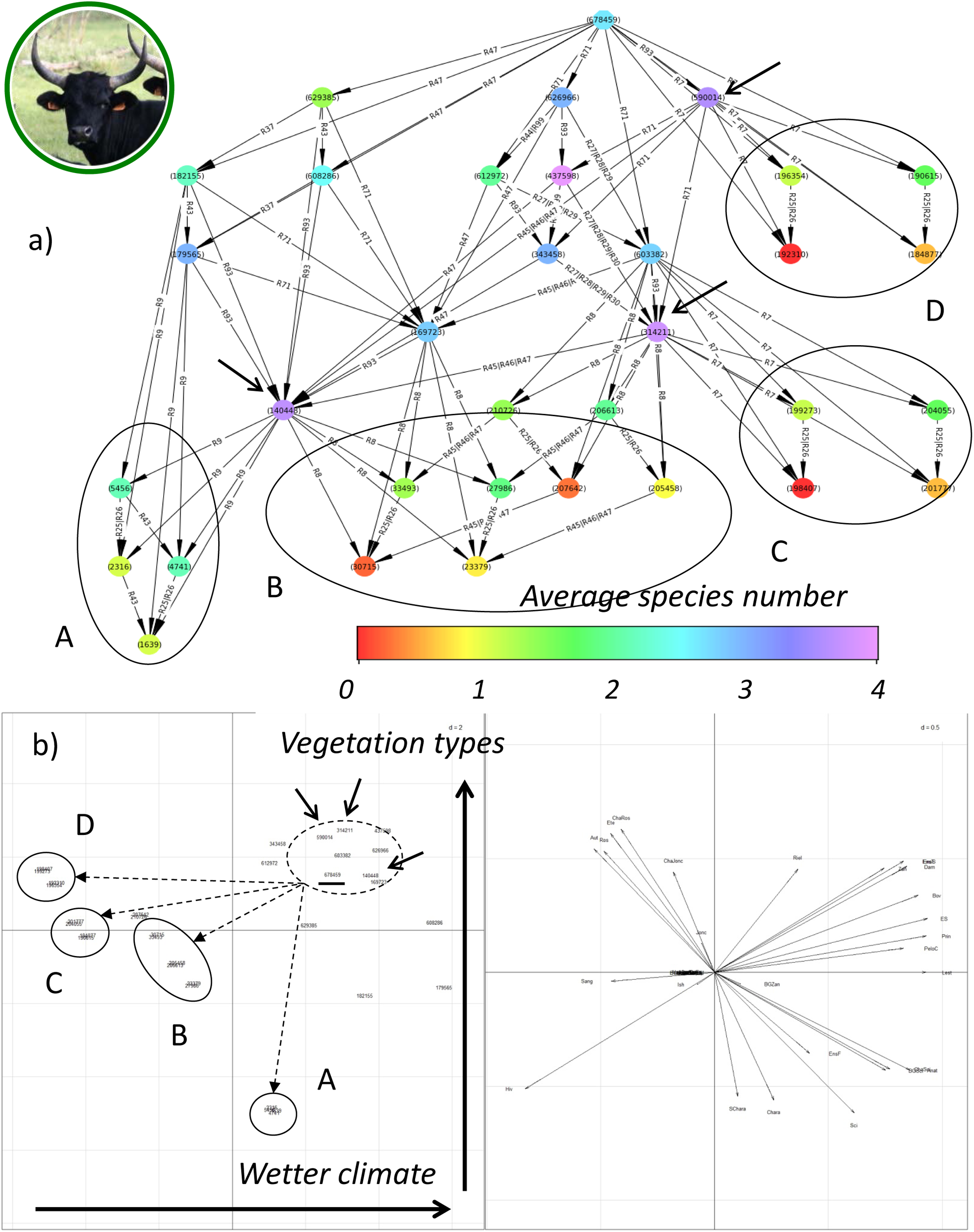
The merged state space of the SBG (seasonal bovine grazing) scenario (a), and its corresponding statistical analysis (PCA, b). The merged state space should be read from top to bottom, through the various trajectories connecting every strongly connected component (a, disks) with various rules (on arrows). The components are gathered (a, ellipses) according to their locations in the PC1/PC2 plan of the PCA (b, left), and each component is colored according to its location on PC1. The statistical analysis (b, right) is easily interpreted according to the projections of the ecosystem nodes involved in each component, and allows reconstructing the ecosystem trajectories (a, ellipses and dashed arrows) in terms of ecosystem composition (b, dashed arrows)..

The trajectories projected by the model, notably the long-term effects of extensive bovine grazing, would be almost impossible to obtain through long-term experiments or by more traditional modeling method. Additionally, the results of the model demonstrate which negative transitions can unlikely be reversed and at what early stage these transitions may be avoided by specific changes in either the management of the herd or the system hydrodynamics.

## Discussion

This study is a step towards a more integrated understanding of any complex socio-ecosystem as a whole, simultaneously to a long term trajectory analysis. It avoids excluding any ecosystem component or process type from the conceptual framework proposed here (Gaucherel et al. 2019), with a system simplification taking place only in later stages of the model building. The interaction network (graph) representation, combined to a topological handling of it, allows reaching this genericity in ecosystem integration. The size and complexity of the model remains feasible mainly due to its fully qualitative conception, thus allowing a generic but rigorous formalization of the *ecosystem development* (Gaucherel et al. 2017). Similar models have been developed in computer sciences and in biology, for example on molecular signalling networks (Blätke et al. 2011) but, to our knowledge, they have never been applied in ecology. Other qualitative models were used for ecological studies, yet focused on small and rather unrealistic interaction networks (Dambacher et al. 2003; Gaucherel and Pommereau 2019).

### Extensive grazing favours temporary marsh biodiversity

In this paper, we illustrated the integrated model with a Petri net approach in the specific case study of temporary Mediterranean marshes in the Camargue. Temporary marshes studied here are protected habitats located in a wetland landscape and hosting several heritage species (Beltrame et al. 2013; Grillas and Roché 1997). Temporary Mediterranean marshes are recognised as a vulnerable habitat in the European Habitat Directive and, as such, form the direct object of conservation management plans. One of the means that is often included in conservation management plans to maintain this habitat includes extensive grazing, but this management entails costs as in most economical-ecological balance (Cote and Nightingale 2012; Cumming et al. 2014). Comparing the results from scenarios without and with extensive grazing confirms and even demonstrates the hypothesis that extensively grazed marshes tend to on average host more species with conservation value (Fig 4a & 5a). This observation is in good agreement with the literature (Bouahim et al. 2010; Noy-Meir et al. 1989; Sternberg et al. 2000). Our innovative applied approach also confirms that the increase or decrease of marsh species richness is not systematic, suggesting that is not unequivocally related to abiotic (hydro-climate) and anthropic (grazing) influences (Fig. 4a-5a, varying colored trajectories). Basically, the model suggests that bovines by their trampling and grazing tend to reduce the salinity of the marsh and then open up the vegetation toward lower strata of the vegetation. Whereas, at high stocking rates, monospecific vegetation outcomes are more likely (either rushes, scirpae or reeds, Fig. 5b).

To validate the system development (trajectories) as coherent is a kind of expert *qualitative validation* of the model. Computer specialists apply two separate ways for validating such innovative and qualitative models (Reisig 2013): either the model is deemed *correct* (i.e. some modeled trajectories, at least, are observed or logical), or the model is deemed *complete* (all trajectories modeled are known/observed). Here, we assumed a list of potentially isolated processes, to finally deduce and confirm a known overall (holistic) behaviour of the studied socio-ecosystem. While each process included in the marsh model was validated independently either by literature or experts (Appendix 4, Table S5-S6), their numerous feedbacks and the integrated response of the ecosystem as a whole was not preconceived.

Here, we were interested in a set of species specifically linked to temporary Mediterranean marshes (Appendix 3, Table S2), and we modelled an increase in these number of these species under extensive bovine grazing (i.e. correctness). This observation is in good agreement with what specialists know about temporary marshes (Beltrame et al. 2013; Grillas and Roché 1997; Rhazi et al. 2006), and with the intermediate disturbance theory (Fox 2013; Wilkinson 1999). This theory suggests that species number of temporary marshes would, over the long term, decrease under no or under intense grazing regimes (Duncan 1992; Grillas et al. 2004), due to differential negative impacts of cattle on vegetation. Here, we focused on extensive bovine grazing, but many other impacts on the ecosystem components have been neglected (e.g. invasive species). These trajectories under various grazing pressures are a qualitative validation of our model able to recover them. Although our found ecological results have already observed in previous studies, this paper demonstrates that our innovative model and conception (Gaucherel et al. 2019) is capable to capture and reproduce this well-known behaviour of such complex socio-ecosystems. Given that it covers multiple transitions simultaneously, it becomes a valuable help for management to identify sustainable scenarios by taking into account all relevant components and processes (Beltrame et al. 2013; Mesléard and Perennou 1996). Conversely, such a qualitative model appears quite powerful in the context of few or lacking data and knowledge on the studied socio-ecosystem, as it would require a simplified qualitative assessment on hypothesized processes. Both these perspectives appear at hand.

### Power and drawbacks of discrete models for ecosystem development

An increasing number of integrated ecosystem models are being developed today (e.g. Gaucherel and Pommereau 2019; Ings et al. 2009; Kéfi et al. 2016), and with them the integration of abiotic, biotic and human components is also improving. Although the number of ecosystem models increases (Dambacher et al. 2003; Lafond et al. 2017; Thébault and Fontaine 2010), they usually focus on relatively short term fluxes created by frozen structures and rarely on alternative developmental (structural) trajectories. In addition, tools to handle the component integration and a rigorous (formalized) system framework are still lacking. In this paper, we developed a discrete, qualitative and non-stochastic model, based on Petri nets (Pommereau 2010; Reisig 2013), to analyse system properties and long term changes (fates). These kinds of models can potentially be extended into quantitative, spatially explicit and multi-temporal schemes to allow more realistic and applied analyses. The real strength of Petri net models is to exhaustively explore all possible trajectories. Indeed, to focus on qualitative (structural) changes of the system or on topological changes of the whole interaction network is equivalent to explore the system changes over a long time period (Gaucherel and Pommereau 2019; Gaucherel et al. 2017).

These early Petri nets of ecosystems are in a preliminary stage and still require a number of improvements. For example, as in every modelling task, the model definition remains partly subjective and dependent on the network definition (Blätke et al. 2011; Gaucherel and Pommereau 2019). In ongoing works, we are testing strict protocols for model building and analyses, for instance in the characterization of nodes (Appendix 3). These protocols increase the objectivity in the choice of nodes and processes when defining the ecosystem model. Also, sensitivity of the model may vary depending on individual components and processes. More research is required to develop sensitivity tests and, thus, a deep understanding of the impact of this variability on its emergent properties. Also, the PCA statistics used here help analyzing the computed state space, but are not yet an interpretation tool in such possibilistic space. Finally, the calibration and validation stages remain real challenges for such integrated and exhaustive (Possibilistic) models. Developing a qualitative rather than a quantitative model significantly reduces the amount of data that needs to be collected for the parameterisation: the model is an abstraction and remains simple. The cost of this simplicity is the model validation, often limited to expert knowledge on known ecosystem trajectories (Gaucherel and Pommereau 2019). As in any model, unknown trajectories cannot be validated nor easily dismissed.

The demonstrated model approach simplifies interannual interactions and processes (i.e. ecosystem functioning), and focuses on long term and structural changes in the ecosystem trajectories. Since the famous Clements/Tansley debate about the organismic nature of any (socio-)ecosystem (Gaucherel 2014; Gaucherel et al. 2019; Gignoux et al. 2011), related concepts to understand ecosystem dynamics have been regularly proposed. These include concepts such as vegetal successions, regime shifts or resilience (Holling 1973; Walker et al. 2004), but these generally focus on one single component of a complex system (e.g. the vegetation) and/or one type of changes (e.g. temperature shifts). The demonstrated modelling approach is one of the few concepts that can grasp all interactions that underlie the structural and qualitative changes at a system level, whatever the ecological or human interactions (Geijzendorffer et al. 2017; Ostrom 2009), and we identified here all the components and processes responsible for specific responses of the integrated socio-ecosystem modelled.

## Acknowledgment

We thank the INRA ECOSERV Meta-program for his financial support. The contribution of IGR was partly funded by ECOPOTENTIAL project (Contract No. 641762) funded under the Horizon 2020 Programme of the European Commission. We thank the experts at the Tour du Valat for their open exchanges, in particular Damien Cohez, Lisa Ernoul, Hugo Fontes, Patrick Grillas, Philippe Lambret, François Mesleard, Anthony Olivier and Olivier Pineau.

## Appendices

Appendix 1: Temporary marsh functioning

Appendix 2: Formal definition of the march model

Appendix 3: Protocol for the model definition

Appendix 4: Rules and constraints of the temporary marsh model

## References

Baldan, P., M. Bocci, D. Brigolin, N. Cocco, and M. Simeoni. 2015, Petri nets for modelling and analysing trophic networks. M. Heiner, and A. K. Wagler, eds. BioPPN 2015, a satellite event of PETRI NETS 2015 1373.

Beltrame, C., E. Cohen-Shacham, M. Trouillet, and F. Guillet. 2013. Exploring the links between local management and conservation applying the ecosystem services concept: conservation and tourism service in Camargue, France. International Journal of Biodiversity Science, Ecosystem Services & Management 9:166–177.

Blätke, M. A., M. Heiner, and W. Marwan. 2011, Tutorial. Petri Nets in Systems Biology. Magdeburg, Germany, Otto-von-Guericke University

Bouahim, S., L. Rhazi, B. Amami, N. Sahib, M. Rhazi, A. Waterkeyn, A. Zouahri et al. 2010. Impact of Grazing on the Species Richness of Plant Communities in Mediterranean Temporary Pools (Western Morocco). Comptes Rendus Biologies 333:670–679.

Brose, U., R. J. Williams, and N. D. Martinez. 2006. Allometric scaling enhances stability in complex food webs. Ecology Letters 9:1228–1236.

Campbell, C., S. Yang, R. Albert, and K. Sheab. 2011. A network model for plant-pollinator community assembly. Proceedings of the National Academy of Sciences (PNAS) 108:197–202.

Chambers, P. A., and E. E. Prepas. 1990. Competition and coexistence in submerged aquatic plant communities: the effects of species interaction versus abiotic factors. Fresh. Biol. 23:541–550.

Cincotta, R. P., J. Wisnewski, and R. Engelman. 2000. Human population in the biodiversity hotspots. Nature 404:990–992.

Cote, M., and A. J. Nightingale. 2012. Resilience thinking meets social theory : Situating social change in socio-ecological systems (SES) research. Progress in Human Geography 36:475–489.

Cumming, G. S., A. Buerkert, E. M. Hoffmann, E. Schlecht, S. v. Cramon-Taubadel, and T. Tscharntke. 2014. Implications of agricultural transitions and urbanization for ecosystem services. Nature 515 50–57.

Dambacher, J. M., H.-K. Luh, H. W. Li, and P. A. Rossignol. 2003. Qualitative Stability and Ambiguity in Model Ecosystems. The american naturalist 161:876–888.

Duncan, P. 1992. Equids in Grazing Systems, Pages 195-204 Horses and grasses. The Nutritional Ecology of Equids and Their Impact on the Camargue, Springer.

Fox, J. W. 2013. The intermediate disturbance hypothesis should be abandoned. Trends in Ecology & Evolution 28:86–92.

Frontier, S. P.-V. D., A. Lepêtre, D. Davoult, and C. Luczak. 2008, Ecosystèmes. Structure, Fonctionnement, Evolution: Sciences SUP. Paris, Dunod.

Gaucherel, C. 2014. Ecosystem Complexity through the Lens of Logical Depth: Capturing Ecosystem Individuality. Biological Theory:10.1007/s13752-13014-10162-13752.

Gaucherel, C. 2019. Physical concepts and ecosystem ecology: a revival? Journal of Ecosystem and Ecography In press.

Gaucherel, C., F. Boudon, T. Houet, M. Castets, and C. Godin. 2012. Understanding Patchy Landscape Dynamics: Towards a Landscape Language. PLoS One 7:e46064.

Gaucherel, C., P. H. Gouyon, and J. L. Dessalles. 2019, Information, the hidden side of life: Information systems, web and pervasive computing series. London, UK, ISTE, Wiley.

Gaucherel, C., and F. Pommereau. 2019. Using discrete systems to exhaustively characterize the dynamics of an integrated ecosystem. Methods in Ecology and Evolution In press.

Gaucherel, C., H. Théro, A. Puiseux, and V. Bonhomme. 2017. Understand ecosystem regime shifts by modelling ecosystem development using Boolean networks. Ecological Complexity 31:104–114.

Geijzendorffer, I., E. Cohen-Shacham, A. F. Cord, W. Cramer, C. Guerra, and B. Martín-López. 2017. Ecosystem services in global sustainability policies. Environmental Science & Policy 74:40–48.

Giavitto, J. L., and O. Michel. 2003. Modeling the topological organization of cellular processes. Biosystems 70:149–163.

Gignoux, J., I. D. Davies, S. R. Flint, and J.-D. Zucker. 2011. The ecosystem in practice : interest and problems of an old definition for constructing ecological models. Ecosystems 14 1039–1054.

Gough, L., and J. B. Grace. 1998. Herbivore effects on plant species density at varying productivity levels. Ecology 79:1586–1594.

Grillas, P., P. Gauthier, N. Yavercovski, and C. Perennou. 2004. Mediterranean Temporary Pools. Arles (France), Station Biologique de la Tour du Valat.

Grillas, P., and J. Roché. 1997, Vegetation of temporary marshes. Ecology and management: Conservation of Mediterranean wetlands. Arles (France), Station Biologique de la Tour du Valat. ed.

Holling, C. S. 1973. Resilience and stability of ecological systems. Annu. Rev. Ecol. Syst. 4:1–23.

Ings, T. C., J. M. Montoya, J. Bascompte, N. Blüthgen, L. Brown, C. F. Dormann, F. Edwards et al. 2009. Ecological networks – beyond food webs. Journal of Animal Ecology 78:253–269.

Kéfi, S., V. Miele, E. Wieters, S. Navarrete, and E. Berlow. 2016. How Structured Is the Entangled Bank? The Surprisingly Simple Organization of Multiplex Ecological Networks Leads to Increased Persistence and Resilience. PLoS Biology 14:e1002527.

Lafond, V., T. Cordonnier, M. Zhun, and B. Courbaud. 2017. Trade-offs and synergies between ecosystem services in unevenaged mountain forests: evidences using Pareto fronts. European Journal of Forest Research:10.1007/s10342-10016-11022-10343.

Lefever, R., and O. Lejeune. 1997. On the origin of tiger bush. Bulletin of Mathematical Biology 59:263–294.

Medail, F., H. Michaud, J. Molina, G. Paradis, and R. Loisel. 1998. Conservation de la flore et de la végétation des mares temporaires dulcaquicoles et oligotrophes de France méditerranéenne. Ecol. Medit. 24:119–134.

Mesléard, F., and C. Perennou. 1996, Aquatic emergent vegetation. Ecology and management: Conservation of Mediterranean wetlands. Arles (France), Station Biologique de la Tour du Valat. ed.

Noy-Meir, M. Gutman, and Y. Kaplan. 1989. Responses of Mediterranean grassland plants to grazing and protection. J. Ecol. 77:290–310.

Ostrom, E. 2009. A general framework for analyzing sustainability of social-ecological systems. Science 325:419–422.

Pommereau, F. 2010. Algebras of coloured Petri nets. Lambert Academic Publishing (LAP). Pommereau, F. 2015, SNAKES. A flexible high-level Petri nets library Proceedings of PETRI NETS’15. LNCS 9115.

Reisig, W. 2013, Understanding Petri Nets. Berlin, Heidelberg, Springer Berlin Heidelberg.

Rhazi, L., M. Rhazi, P. Grillas, and D. El Khyari. 2006. Richness and structure of plant communities in temporary pools from western Morocco: influence of human activities. Hydrobiologia 570:197–203.

Ricklefs, R. E., and G. L. Miller. 2000, Ecology. New York, USA, Freeman (4th edition).

Sternberg, M., M. Gutman, A. Perevolotsky, E. D. Ungar, and J. Kigrl. 2000. Vegetation response to grazing management in a Mediterranean herbaceous community: a functional group approach. J. App. Ecol. 37:224–237.

Thébault, E., and C. Fontaine. 2010. Stability of Ecological Communities and the Architecture of Mutualistic and Trophic Networks. Science 329:853–856.

Titeux, N., K. Henle, J. B. Mihoub, A. Regos, I. R. Geijzendorffer, W. Cramer, P. H. Verburg et al. 2017. Global scenarios for biodiversity need to better integrate climate and land use change. Diversity and Distributions 23:1231–1234.

Walker, B., C. S. Holling, S. R. Carpenter, and A. Kinzig. 2004. Resilience, adaptability and transformability in social-ecological systems. Ecology and Society 9:5.

Wilkinson, D. M. 1999. The Disturbing History of Intermediate Disturbance. Oikos 84 145–147.

