## Appendices 1 to 4 for "Long term development of a realistic and integrated socio-ecological system"

**Development in the long term of an integrated and dynamical ecosystem**

**Gaucherel C. ^(1) *^, Carpentier C. ^(1, 2)^ Geizjendorffer I. ^(3)^ Pommereau F. ^(4)^**

### SUPPLEMENTARY MATERIALS - APPENDICES

### Appendix 1: Temporary marsh functioning

#### Temporary marshes

Temporary Mediterranean pools are small shallow depressions (<10 ha) fed by precipitation in autumn or spring. These pools are characterized by the alternation of flooding and drying out phases (Grillas et al., 2004a). The succession of these contrasting ecological phases provides a niche for many aquatic, amphibian and terrestrial species. Indeed, these pools are submerged during sufficiently long intervals of time and, thus, allow the development of a flora rich in aquatic (*e.g. Zannichellia sp.)* and amphibious (*e.g. Damasonium polyspermum*) species (Grillas and Roché, 1997). These species are often totally dependent on temporary pools: uncompetitive, they have developed various adaptations to resist the stressing seasons. Hence, species observed on these sites are specific and ubiquist species, while those of permanent waters cannot settle (Grillas et al., 2004a).

The species of these environments usually have life cycles with two phases, aquatic and terrestrial ones. Consequently, temporary pools provide habitats for remarkable amphibians (*e.g. Pelobates cultipes*) or dragonflies (*e.g. Lestes macrostigma*) (Grillas et al., 2004b). Conversely, ichthyofauna is absent from these environments: since connections to permanent waters are rare, few fish reach temporary pools and even less escape before pools dry out (Grillas and Roché, 1997). Moreover, the abundance of plant and animal resources in these environments makes it a feeding habitat for many birds (*e.g. Anas crecca*), but also for wild mammals (*e.g. Sus scrofa*) and domestic ones. The latter graze pools during their terrestrial phase (*e.g.* ovines and caprines) and / or their drying out phase (*e.g.* equines and bovines). The presence of such animals entail various disturbances such as trampling, browsing, and fertilizing inputs (Mesléard and Perennou, 1996).

#### Grazing impacts and scenarios

At the Tour du Valat, the temporary pools account for 250 ha (Cohez et al., 2016). Only a small number of them are recognized as brackish "lagoons" and therefore this behaviour will not be modelled. Moreover, the model constructed describes temporary pools in the current context in which they occur: dyked Rhône, controlled hydrological network, absence of *Oryctolagus cuniculus*, etc. Finally, the model considers a pool dynamics without any other major disturbance than the grazing by domestic animals (*i.e.* no invasive species, no connection with permanent water, etc.).

The impact of grazing on temporary pools flagship species is determined by comparing different scenarios. These scenarios are simulated on the basis of a unique set of nodes and a unique set of rules: only the initial state differs (see Appendix 1). This allows some rules to be enabled in some scenario but not in other. Because the enabled rules set changes, a unique state space is associated with each scenario. However, since these scenarios are based on the same nodes and rules, it is quite likely that some of their states are in common – except for the node modified in the initial state. In this context, ignoring these nodes allow finding identical states in the different scenarios. These states correspond to state where scenarios do not differ. Moreover, some states will not be in common between scenarios: such states inform on the impact of each scenario.

### Appendix 2: Formal definition of the marsh model

#### Formal model: nodes, rules, and state space

A detailed description of Petri net models may be found in the literature (Reisig 2013, Gaucherel and Pommereau 2019). We give here the minimal description allowing reproducing our results. In particular, we omit the definition of Petri nets, as well as the transformation from rules to Petri nets.

A system consists of Boolean nodes that represent the states of agents in the ecosystem, as well as of rules and constraints that define how nodes values may evolve applying an effect (assignment of the nodes values) and depending on a guard (condition on the nodes values). Constraints have a higher priority than rules and are used to model cascading events in ecosystems. For instance, if a habitat disappears, all its inhabitants must disappear too. Figure 2b depicts a simple predator-prey system modelled in this way. Nodes are declared as a name, an initial state, and a textual description. For instance, we declare node P that is initially ON (+) and represents the presence of predators in the (eco)system. These declarations are organised into arbitrarily chosen categories (except for “rules” and “constraints” that are reserved keywords), thus belonging to only one category called “components”. Rules and constraints are listed afterward and consist in two-sides separated by “>>”: left-hand side is the guard, that is, a condition for the execution of the rules; right-hand side is the effect, that is, the nodes assignments that results from the execution of the rule. For instance, rule $R_{1}$ specifies that, in the presence of predators, preys may disappear (they are eaten, i.e. their population is reduced to a few individuals having no more impact on the system functioning). Similarly, rule $R_{2}$ specifies that, in the absence of preys, predators may disappear (they die or are functionally absent).

The semantics of such systems has been originally defined both in terms of operational rules, as well as in terms of a translation to Petri nets, both semantics being proved equivalent (Gaucherel and Pommereau 2019). Intuitively, the semantics is as follows:

- the *initial state* is that defined from the declaration of the entities;
- at some state, the *successors* are obtained by executing constraints;
- if no constraint is enabled, then the successors are obtained by executing rules;
- if no rule nor constraint is enabled, then the system is in a *deadlock*, a state from which the system does not evolve anymore;
- an *action*, which is a rule or a constraint, is enabled when its guard is validated by the state and its effect is not already realised (i.e. we forbid executions that do not change the state);
- executing an action $a$ enabled at a state $s$ is made by applying the effect onto $a$, yielding a new state $s^{'}\neq s$, which is called a *transition* noted by $s \underset{\to}{a}s^{'}$.

For instance, the model defined in Figure 2b has the initial state $\{P,N\}$ (we just list the nodes that are ON). From this state, rule $R_{2}$ is not enabled because its condition is not validated; but rule $R_{1}$is enabled because we have $P=ON$ and not yet $N=OFF$. So we may execute $R_{1}$yielding a new state $\{P\}$. This is represented in Fig 2c by the arc $S_{0}\underset{\to}{R_{1}}S_{1}$.

Given a model, its *state space* is a triple $(s_{0}, S,A,\to)$, where $s_{0}$ is the initial state, and $S$, $A$, and $\to$ are the smallest sets such that $s_{0}\in S$, and, whenever $s\underset{\to}{a}s^{'}$ for some $s\in S$, we also have $s^{'}\in S$, $a\in A$, and $\left( s,a,s^{'} \right)\in\to$.

#### Compact state space

Constraints are made up of a condition and a realisation, just as rules do, and represent unavoidable events given the system state (which is implemented by their priorities over rules).

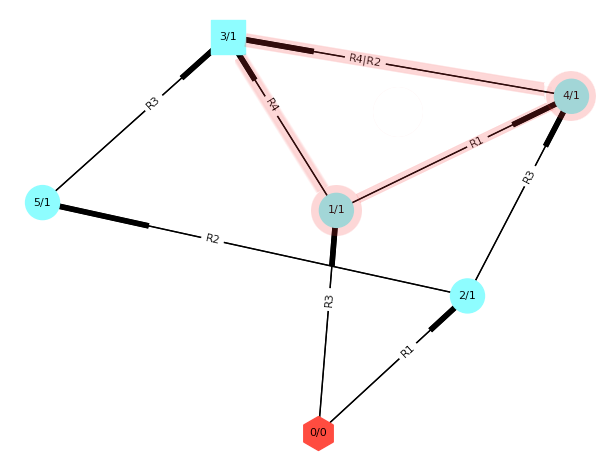

Rules used

R1: PF+ >> IF-

R2: IF- >> PF-

R3: S+ >> P-

R4: P- >> PF-, IF-

Figure S1. Nodes and rules defining the pond-fishes system and the State space associated with the model described in the table and showing only present nodes, s_0_={PF, IF, S, P}; s_1_={PF, IF, S}; s_2_={PF, S, P}; s_3_={S}; s_4_={PF, S}; s_5_={S, P}. Unrealistic paths are overwritten in red.

As an example, we define a shallow pond (denoted P) that may dry out in summer (S) and contain insectivorous (IF) and piscivorous fishes (PF). Such a system may be described by four nodes and four rules resulting in a state space of size six (Fig. S1). In this state space, the application of rule R3 leads to a realistic state (s_3_) and two unrealistic ones (s_1_ and s_4_). In the latter states, fishes continue to live and interact while no more water is in the pond. Indeed, rule R4 (associated with the disappearance of fishes when water dries out) is considered as a “possibility” after R3 firing and not as an unavoidable event. For this reason, R4 should be a constraint: it *must* be applied, otherwise the model simulates unrealistic paths.

The *compact state space* is defined to keep only realistic states while dropping transient states. Let $s$ and $s'$ be two realistic states we note by $s\underset{\to}{R\cdots}s^{'}$ the fact that $s\underset{\to}{R}s'$ for some rule $R$ or $s\underset{\to}{R}s^{''}\underset{\to}{C_{1}}\cdots\underset{\to}{C_{k}}s'$ for some rule $R$, an unrealistic state $s^{''}$, and constraints $C_{1}$ to $C_{k}$ ($k\geq1$), i.e., $s^{'}$ is reachable from $s$ through exactly one execution of a rule $R$, possibly followed by the executions of some constraints. We extend this notation to pairs of sets like $s^{''}$ and $s'$ above where a realistic state $s^{'}$ can be reached from an unrealistic $s''$ through constraints only. With this notation, the compact state space is defined as a triple $(S_{0}, S_{r},A,\underset{\to}{\ldots})$, where: $S_{0}$ is either $\{s_{0}\}$ if $s_{0}$ is realistic, or $S_{0}=\{s\mid s_{0}\underset{\to}{R\ldots}s\}$ otherwise; $S_{r}\subseteq S$ is the set of realistic states; and $\underset{\to}{\ldots} =\{\left( s,R,s^{'} \right)\mid s,s^{'}\in S_{r}\wedge s\underset{\to}{R\cdots}s^{'}\}$. Using constraints instead of rules allows an efficient reduction of state space size in the temporary marsh models (Table S1).

Table S1. Number of all possible states and realistic states only in the different models of this study.

|  | | # Possible states | # Realistic states | Ratio |
| --- | --- | --- | --- | --- |
| Models | Marsh model | 1.60 x 10^1^ | 0.40 x 10^1^ | 2.50 x 10^-1^ |
|  | Marsh without grazing | 7.03 x 10^13^ | 1.60 x 10^6^ | 2.27 x 10^-8^ |
|  | marsh with grazing | 7.03 x 10^13^ | 1.54 x 10^6^ | 2.18 x 10^-8^ |

#### Merged state space

As presented above, a state space $(s_{0},S,A,\to)$ (or a compact state space $\left( S_{0}, S_{r},A,\underset{\to}{\ldots} \right)$, this makes no difference) can be seen as a graph, whose vertices are $S$ and whose edges are $\to$, with labels from $A$. In such a graph, one can distinguish sets of vertices/states:

- the *initial state* is $s_{0}$;
- the *strongly connected components* (SCC) are maximal subsets of states such that there is a path between any pair of nodes in a SCC. Possibly, the initial state is included in a SCC;
- the *deadlocks* are states from which no other state is reachable (i.e. a structural dead-end);
- altogether, these sets are called the *primitive components*;
- the *basins of attractions* are maximal subsets of states that all allow to reach exactly the same primitive components. A basin of attraction that leads to only a deadlock may be merged with this deadlock to reduce the number of components.

These components form a partition of the states and allow to define a merged state space as the smallest graph $\left( S_{0},C,A,↪ \right)$ such that: $S_{0}$ is either the SCC that contains $s_{0}$ if any or the singleton $\left\{ s_{0} \right\}$ otherwise; $C$ is the set of components as defined above; whenever there exists $s\in c\in C$ and $s^{'}\in c^{'}\in C$ such that $s \underset{\to}{a}s'$, then we have $\left( c,a,c^{'} \right)\in↪$.

The benefits from considering this merged state space is that it is much smaller than the original state space while still exhibiting the important dynamics of the system. However, in the case of the Camargue model, the merged state space is still very large because it has a lot of small size SCCs. Indeed, the rules modelling the cycle of seasons and rain may be executed alone, yielding to a small SCC with a single state (that would otherwise be classified in a basin of attraction). In order to get rid of these small SCCs that do not convey relevant information, we have deleted them from the merged graph. Doing so, we lose some states and some trajectories. In order to ensure that we did not remove existing behaviours in the systems, we checked that for every pair of components reachable one from the other, this reachability is preserved when we remove the small SCCs. In other words, we have removed some trajectories but there still exists equivalent ones as far as reachability is concerned.

### Appendix 3: Protocol for the model definition

#### Node characterization

We highlight three levels of *characterization* for a node:

- The node is *fully* *characterized* if at least one rule or constraint sets it ON and at least one rule sets it OFF;
- The node is *semi***-***characterized* if it can never return to its previous state, i.e., it is initially ON and at least one rule or constraint sets if OFF but no rule is able to set it ON again, and the other way round;
- The node is *fixed* if its state cannot vary in the system, i.e., it does not appear in the right-hand side of a rule or constraint with a value that is the opposite of its initial state.

In an ideal model, most nodes would be fully characterized, as most of them in real life influence and are influenced. Hence, semi-characterized nodes must be merged with others or removed from the graph and the rules have then to be modified to take into account for this rearrangement of the ecosystemic graph. Some nodes may be fixed if they have an impact on the defined ecosystem, but are not influenced by it (*e.g.* summer in the pound model, Appendix 4). Finally, the characterisation degree of a node is defined as the number of rules in the right-hand side of which it appears.

We may distinguish the *static characterization* as defined above from the *dynamic characterization* which is based on the actual executions of rules. Indeed a node may be fully characterized by some rules that are actually never executed so that the node turns out to be fixed. So, dynamic characterization can be defined exactly as above but considering only the rules and constraints that are actually executed in the state space.

The node characterization is one of the most important stages in the model building procedure. It defines the set of nodes and the set of rules describing the studied ecosystem. Consequently, choices made by the modeller at this stage have consequences on all the following steps, from the graph folding to the trajectory analysis (see Appendix 2). The state of a node varies more if more rules impact this node. Hence, the *characterization degree* is determined by two criteria: the ecology of the given node and the knowledge available to the modeller. We must maximize the influence of the first criterion and minimize the latter. The modeller knowledge should indeed not influence the probability that a node switches from one state to another and the distribution of the characterization degree should thus follow a uniform law. However, degrees observed in ecosystems most often follow other distributions (*e.g.* power law for trophic networks (Proulx et al. 2005)). For example in our model, *Scirpus maritimus* is grazed by wild and domestic animals and is competing for light with *Phragmites australis* and wilts in case of long drought.

The variation of a node state within the state space also depends on the length of the condition part of the rules that influence it: the more a rule has nodes in its condition, the less often it is available. Hence, the precision with which the rules are written influences their probability of being fired and, thus, the state space size. Furthermore, semi-characterized nodes impact the state space, by opening one-way paths and thus, fragmenting it. Such nodes seem to be essential to the system dynamics, whereas this emergence is not due to their role in the ecosystem rather than to a lack of knowledge, to modeller negligence, or both. Hence, full characterization of all nodes is essential to model a coherent system dynamics, unless the user consciously wants to split the state space (e.g. invasive species that the system could not remove anymore). In the same logic, fixed nodes also allow limiting the system combinatory: only their initial state impacts the ecosystem trajectories. Therefore, such nodes are used to make processes explicit (*e.g.* “wind” node in the temporary marsh model, Appendix 4) and / or to explore different scenarios based on the same model (*e.g.* the nodes associated to grazing scenarios).

#### Heritage species

In our model, we focused on: i) the hydrology of a temporary marsh, ii) the requirements of domestic mammals, and iii) flagship species characteristic of these environments. For each topic, we identified nodes of interest based on literature and expert informal interviews: surface water for temporary marshes, bovines and equines for domestic mammals and species with patrimonial issues listed by the Tour du Valat (Cohez et al. 2016). The requirements of each component led to a prototypal model (noted Mod1) made up of 112 nodes (including 52 semi-characterized and 10 fixed nodes), 223 rules *stricto sensu* and 92 constraints (see Appendix 2 for formal definitions). We then built a new and condensed version of the model (Mod2) in which 27 of the Mod1 semi-characterized nodes became fully characterized, 22 nodes were deleted or merged with other nodes, and three fixed nodes were ultimately deleted. The model Mod2 thus comprises 87 nodes, 164 rules *stricto sensu* and 123 constraints. This model was finally *folded* into 46 nodes, 105 rules *stricto sensu* and 57 constraints (Appendix 4). This final Mod3 model gathers 17 nodes of interest, among which the heritage species (Table S2) and 29 nodes mandatory for the ecosystem functioning (Table 1).

Table S2. Nodes of interest (heritage species) in the final temporary marsh model.

| Surface water  (Key of the pool dynamics) | Seasonal equine, bovine and ovine grazing  (Rare) |
| --- | --- |
| Bovines  (Herd of cultural heritage) | *Damasonium sp.*  (Alismata of TdV major responsibility) |
| Equines  (Herd of cultural heritage) | *Pelobates cultripes*  (Anura of TdV strong responsibility) |
| Ovines  (Herd historically used in Camargue) | *Lestes macrostigma*  (Odonata of TdV strong responsibility) |
| Caprines  (Herd never used in Camargue) | *Zannichellia sp.*  (Alismata of TdV strong responsibility) |
| Intensive bovine grazing  (Farmers strategy) | *Characeae*  (Charales of TdV quite strong responsibility) |
| Intensive equine grazing  (Farmers strategy) | *Riella helicophylla*  (Marchantiopsida of TdV quite strong responsibility**)** |
| Seasonal bovine grazing  (TdV strategy) | *Isnhura pumilio*  (Odonata used as indicator, specific species) |

The most characterised nodes are those associated with the question addressed (*i.e.* species grazed by domestic animals and flagship species of temporary marshes, Fig. S2). Nodes relative to the atmosphere are the least characterized. The wind and nodes related to the grazing scenario are fixed.

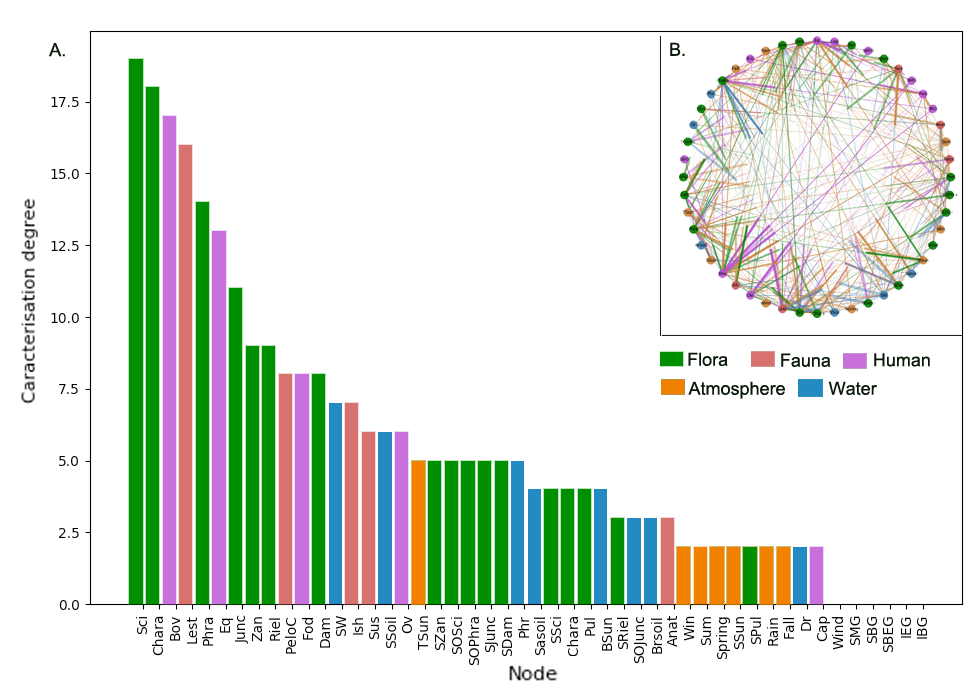

Figure S2. Characterisation degree distribution of nodes in the final model (Mod3), and insert showing the ecosystemic graph associated to it.

### Appendix 4: Rules and constraints of the temporary marsh model

Table S5. Model processes and rule categories, identification number, associated formulae and descriptions of the temporary marsh ecosystem graph. The source of each rule is mentioned: [1] Common sense; [2] (Grillas et al. 2004) ; [3] (Grillas and Roché 1997); [4] (Lambret 2016); [5] (Chazel and Chazel 2013); [6] (Mouronval et al. 2014); [7] (Marnotte et al. 2006); [8] (Mesléard and Perennou 1996); [9] (Willm et al. 2012).

| N° | Rules | Underlying process | |
| --- | --- | --- | --- |
| Atmosphere | | | |
| 1 | Spring+ >> Sum+, Spring- | Seasons cycle | [1] |
| 2 | Sum+ >> Fall+, Sum- | Seasons cycle | [1] |
| 3 | Fall+ >> Win+, Fall- | Seasons cycle | [1] |
| 4 | Win+ >> Spring+, Win- | Seasons cycle | [1] |
| 5 | Rain+ >> Rain- | Sunshine after rain | [1] |
| 6 | Rain- >> Rain+ | Rain after sunshine | [1] |
| 7 | Spring+, SW+,Phra+, TSun+, Sci-, Junc- >> TSun-, BSun- | Dense monospecific cover reduces luminosity | [1] |
| 8 | Sum+, TSun+, SSoil+, Junc+, Phra-, Sci- >> TSun-, BSun- | Dense monospecific cover reduces luminosity | [1] |
| 9 | Sum+, TSun+, Sci+, Phra-, Junc->> TSun-, BSun- | Dense monospecific cover reduces luminosity | [1] |
| 10 | Zan+ >> SSun- | Hydrophytes generate shade | [1] |
| Fauna | | | |
| 11 | Win+, Rain+, SW+ >> PeloC+ | Reproduction of Plobates cultripes takes place during winter rains | [2] |
| 12 | Spring+, Rain+, SW+ >> PeloC+ | Reproduction of Plobates cultripes takes place during spring rains | [2] |
| 13 | Sum+, Lest+ >> PeloC- | Dragonfly larvae predate Plobates cultripes larvae | [2] |
| 14 | Sum+, Ish+ >> PeloC- | Dragonfly larvae predate Plobates cultripes larvae | [2] |
| 15 | Spring+, Sci+, SW+ >> Lest+ | Durign spring, Lestes macrostigma lays eggs in the stems of Scirpus maritimus | [4] |
| 16 | Spring+, Sci-, Junc+, SW+ >> Lest+ | Durign spring, Lestes macrostigma lays eggs in the stems of Juncus sp. if there is no Scirpus maritimus | [4] |
| 17 | Sum+, Sci-, Junc- >> Lest- | If Scirpus maritimus and Juncus sp. disappear before the emergence of the larvae, Lestes macrostigma population is not maintained | [4] |
| 18 | Fall+, Sci-, Junc- >> Lest- | If Scirpus maritimus and Juncus sp. disappear before the emergence of the larvae, Lestes macrostigma population is not maintained | [4] |
| 19 | Win+, Sci-, Junc- >> Lest- | If Scirpus maritimus and Juncus sp. disappear before the emergence of the larvae, Lestes macrostigma population is not maintained | [4] |
| 20 | Win+, SW- >> Lest- | A too late watering of the pool provokes the death of Lestes macrostigma larvae | [4] |
| 21 | Fall+, Junc+, BSun+, SW+ >> Ish+ | During fall, Isnhura pumilio lays eggs in the stems of Juncus sp. (first generation) | [2] |
| 22 | Fall+, Sci+, BSun+, SW+ >> Ish+ | Durign fall, Isnhura pumilio lays eggs in the stems of Scirpus maritimus (first generation) | [2] |
| 23 | Spring+, Junc+, BSun+, SW+ >> Ish+ | During spring, Isnhura pumilio lays eggs in the stems of Juncus sp.(second generation) | [2] |
| 24 | Spring+, Sci+, BSun+, SW+ >> Ish+ | During spring, Isnhura pumilio lays eggs in the stems of bulrush (second generation) | [2] |
| 25 | Rain+ >> Ish- | Isnhura pumilio is a brittle species | [2] |
| 26 | Spring+, SW- >> Ish- | Death of Isnhura pumilio larvae and eggs if spring drying is too long | [2] |
| 27 | Sci+, SW- >> Sus+ | Sus scrofa is attracted by the resources of the pool | [3] |
| 28 | Phra+, SW- >> Sus+ | Sus scrofa is attracted by the resources of the pool | [3] |
| 29 | Junc+, SW- >> Sus+ | Sus scrofa is attracted by the resources of the pool | [3] |
| 30 | Dam+, Riel+, SW- >> Sus+ | Sus scrofa is attracted by the resources of the pool | [3] |
| 31 | Fod+, SW- >> Sus+ | Sus scrofa is attracted by the resources of the pool | [3] |
| 32 | Win+ >> Anat+ | Anatidae winter in the Camargue | [5] |
| Flora | | | |
| 33 | Anat+, SSci- >> Chara- | Anatidae eat Characeae | [6] |
| 34 | Sum+, SSun+, Chara+ >> SChara+ | Chara sp. produces spores in summer | [7] |
| 35 | Fall+, SW+, SSun+, SChara+ >> Chara+ | Chara sp. germinates in autumn if there is enough light | [3] |
| 36 | Spring+, SSun+, Chara+ >> SChara+ | Tolypella sp. produces spores in spring | [7] |
| 37 | Anat+ >> SChara+ | Anatidae participate in the dispersal of Characeae spores | [3] |
| 38 | Anat+, SSci-, SOSci-, Chara- >> SChara- | Grain-eaters Anatidae consume Characeae spores | [6] |
| 39 | Fall+, SW+, BSun+, SZan+ >> Zan+ | Zannichellia sp. germinates in fall if there is enough light | [3] |
| 40 | Spring+, Zan+, BSun+, SW+ >> SZan+ | Zannichellia sp. reproduces in spring if there is enough light | [3] |
| 41 | Fall+, SW+, BSun+, Zan+ >> SZan+ | Zannichellia sp. produces seeds in fall | [3] |
| 42 | Fall+, SW-, SSoil+, Bov+ >> SZan+ | Soil upheaval by animals raises old seeds buried in soil | [3] |
| 43 | Fall+, SW-, SSoil+, Sus+ >> SZan+ | Soil upheaval by animals raises old seeds buried in soil | [3] |
| 44 | Anat+, SSci-, SOSci-, Chara-, SChara- >> SZan- | Grain-eaters Anatidae consume Zannichellia sp. seeds | [6] |
| 45 | Fall+, SW-, SSoil+, SOSci-, Sus+ >> SOPhra-, Lest-, Chara- | In fall, Sus scrofa eats fleshy parts of Phragmites australis and, by turning over the soil, causes the suspension of sediments - which is deleterious for the dragonflies | [10] |
| 46 | Spring+, SW-, SSoil+, SOSci-, Sus+ >> SOPhra-, Lest-, Chara- | In spring, Sus scrofa eats fleshy parts of Phragmites australis and, by turning over the soil, causes the suspension of sediments - which is deleterious for the dragonflies | [10] |
| 47 | Spring+, Bov+ >> Phra-, SOPhra- | In spring, Phragmites australis is pastured by bovines and this grazing prevents re-growth | [8] |
| 48 | Sum+, Bov+ >> Phra- | In summer, Phragmites australis is pastured by bovines | [8] |
| 49 | Spring+, Eq+ >> Phra-, SOPhra- | In spring, Phragmites australis is pastured by equines and this grazing prevents re-growth | [8] |
| 50 | Sum+, Eq+ >> Phra- | In summer, Phragmites australis is pastured by equines | [8] |
| 51 | Spring+, Ov+, Dam-, Riel- >> Phra- | In spring, Phragmites australis is pastured by ovines | [8] |
| 52 | Sum+, Ov+, Dam-, Riel- >> Phra- | In summer, Phragmites australis is pastured by ovines | [8] |
| 53 | Spring+, SW+, SOPhra+, TSun+ >> Phra+ | Phragmites australis grows in summer if there is surface water | [8] |
| 54 | Sum+, SSoil+, Wind+ >> SSci+ | Wind disperses Scirpus maritimus seeds | [8] |
| 55 | Sum+, TSun+, Sci+ >> SSci+ | Scirpus maritimus produces seeds in summer if there is enough light | [8] |
| 56 | Spring+, SOSci+, SW+, TSun+ >> Sci+ | Scirpus maritimus grows in spring if there is surface water and enough light | [8] |
| 57 | Spring+, SSci+, SW+, TSun+ >> Sci+ | Scirpus maritimus germinates in spring if there is surface water and enough light | [8] |
| 58 | Fall+, SW+, Sci+, Phra+ >> Sci- | Late persistence of the water disfavours Scirpus maritimus in competition with Phragmites australis | [8] |
| 59 | Win+, SW+, Sci+, Phra+ >> Sci- | Late persistence of the water disfavours Scirpus maritimus in competition with Phragmites australis | [8] |
| 60 | Fall+, Anat+ >> SSci-, SOSci- | Anatidae feed on Scirpus maritimus seeds and tubers | [6] |
| 61 | Spring+, Anat+ >> SSci-, Sci-,SOSci- | Anatidae feed on Scirpus maritimus stems, seeds and tubers | [6] |
| 62 | Fall+, SW-, SSoil+, Sus+ >> SOSci-, Lest-, Chara- | In fall, Sus scrofa eats fleshy parts of Scirpus maritimus and, by turning over the soil, causes the suspension of sediments - which is deleterious for the dragonflies | [10] |
| 63 | Spring+, SW-, SSoil+, Sus+ >> SOSci-, Lest-, Chara- | In spring, Sus scrofa eats fleshy parts of Scirpus maritimus and, by turning over the soil, causes the suspension of sediments - which is deleterious for the dragonflies | [10] |
| 64 | Spring+, Phra-, Bov+ >> Sci- | Scirpus maritimus is less palatable than Phragmites australis and is pastured by bovines in spring and summer | [8] |
| 65 | Sum+, Phra-, Bov+ >> Sci- | Scirpus maritimus is less palatable than Phragmites australis and is pastured by bovines in spring and summer | [8] |
| 66 | Spring+, Phra-, Eq+ >> Sci- | Scirpus maritimus is less palatable than Phragmites australis and is pastured by equines in spring and summer | [8] |
| 67 | Sum+, Phra-, Eq+ >> Sci- | Scirpus maritimus is less palatable than Phragmites australis and is pastured by equines in spring and summer | [8] |
| 68 | Spring+, Ov+ >> Sci- | Scirpus maritimus by ovines in spring and summer | [8] |
| 69 | Sum+, Ov+ >> Sci- | Scirpus maritimus by ovines in spring and summer | [8] |
| 70 | Sum+, TSun+, SSoil+, Junc+ >> SJunc+ | Juncus sp. produces seeds in summer | [9] |
| 71 | Fall+, TSun+, SW-, SSoil+, SJunc+ >> Junc+ | Juncus sp. germinates in fall if there is water and light | [9] |
| 72 | Fall+, TSun+, SW-, SSoil+, SOJunc+ >> Junc+ | Juncus sp. grows in fall if there is water and light | [9] |
| 73 | Sum+, Anat+ >> SJunc+ | Anatidae participate in the dispersal of Juncus sp. seeds | [9] |
| 74 | Sum+, Sus+, SW-, SSoil+ >> SJunc+ | Sus scrofa participates in the dispersal of Juncus sp. seeds | [9] |
| 75 | Fall+, Sasoil+ >> SJunc- | Juncus sp. germination is affected by high salinity in autumn | [9] |
| 76 | Win+, Anat+ >> SJunc- | Grain-eaters Anatidae eat Juncus sp. seeds | [6] |
| 77 | Bov+, Phra-, Sci- >> Junc- | Juncus sp. is less palatable than Phragmites australis and Scirpus maritimus and is pastured by bovines | [3] |
| 78 | Eq+, Phra-, Sci- >> Junc- | Juncus sp. is less palatable than Phragmites australis and Scirpus maritimus and is pastured by equines | [3] |
| 79 | Cap+ >> Junc-, SOJunc- | Juncus sp. is pastured by caprines | [3] |
| 80 | Sus+, SOPhra-, SOSci-, Fod-, SW- >> SOJunc- | Juncus sp. is not palatable but may be pastured by Sus scrofa | [3] |
| 81 | Bov+, Phra-, Sci-, Junc- >> Dam-, Riel- | Herbaceous species are pastured by bovines in absence of helophytes | [3] |
| 82 | Cap+, Junc- >> Dam-, Riel- | Herbaceous species are pastured by carines in absence of Juncus sp. | [3] |
| 83 | Ov+ >> Dam-, Riel- | Herbaceous species are pastured by ovines | [3] |
| 84 | Spring+, SW+, TSun+, SDam+ >> Dam+ | Damasonium sp. germinates when the pool is under water | [2] |
| 85 | Sum+, Dam+, SW-, SSoil+ >> SDam+ | Damasonium sp. produces seeds in summer if the soil is saturated | [2] |
| 86 | Sum+, Wind+ >> SDam+ | Wind disperses Damasonium sp. seeds | [2] |
| 87 | Sus+, SW- >> SDam-, Lest-, Chara- | Sus scrofa eats Damasonium sp. seeds and, by turning over the soil, causes the suspension of sediments - which is deleterious for the dragonflies | [10] |
| 88 | Bov+ >> SDam- | Bovines trample the soil and prevent Damasonium sp. seed from germination | [2] |
| 89 | Eq+ >> SDam- | Equines trample the soil and prevent Damasonium sp. seed from germination | [2] |
| 90 | Fall+, TSun+, SW+, SRiel+ >> Riel+ | Riella helicophylla germinates in fall if there is enough water and light | [2] |
| 91 | Spring+, SW+, Riel+ >> SRiel+ | Riella helicophylla produces spores in spring if there is enough water | [2] |
| 92 | Spring+, SW+, Anat+ >> SRiel+ | Anatidae participate in the dispersal of Riella helicophylla spores | [2] |
| 93 | Sum+, SSoil+ >> SRiel- | Without drying out in summer, Riella helicophylla spores lose their germinative power | [2] |
| Human | | | |
| 94 | IBG+, Bov+ >> Phra-, Sci+, Zan-, Chara- | A high nitrogen content destroys Phragmites australis, amphibious vegetation, hydrophytes and algae and favors Scirpus maritimus | [10] |
| 95 | IEG+, Eq+ >> Phra-, Sci+, Zan-, Chara- | A high nitrogen content destroys Phragmites australis, amphibious vegetation, hydrophytes and algae and favors Scirpus maritimus | [10] |
| Water | | | |
| 96 | Rain-, Wind+, Sum+ >> SW-, Zan- | Evaporation of surface water is detrimental to hydrophytes | [10] |
| 97 | Rain-, Wind+, Sum+ >> SSoil-, SW-, Chara-, Zan-, Riel-, Dam-, Sci-, Phra-, Junc- | Prolonged drought causes the evaporation of interstitial water - which is harmful to algae, hydrophytes and amphibian species | [10] |
| 98 | Rain-, Wind+, Sum+ >> Phr-, SSoil-, SW-, Brsoil+, Chara-, Zan-, Riel-, Dam-, Sci-, Phra-, Junc-, TSun+ | Prolonged drought causes the evaporation of interstitial water - which is harmful to algae, hydrophytes and amphibian species and causes an increase in salinity | [10] |
| 99 | Rain-, Wind+, Sum+, Brsoil+ >> Phr-, SSoil-, SW-, Sasoil+, Chara-, Zan-, Riel-, Dam-, Sci-, Phra-, Junc-, TSun+ | Prolonged drought causes the evaporation of interstitial water - which is harmful to algae, hydrophytes and amphibian species and causes an increase in salinity | [10] |
| 100 | Rain+ >> Phr+, Brsoil-, Sasoil- | Rains fill the groundwater and leach soil | [3] |
| 101 | Rain+ >> Brsoil-, Sasoil- | Rains leach soil | [3] |
| 102 | Rain+ >> Sasoil- | Rains leach soil | [3] |
| 103 | Rain+ >> Phr+, SSoil+ | Rains fill the groundwater and the soil | [3] |
| 104 | Rain+ >> Phr+, SSoil+, SW+ | Rains fill the groundwater, the soil and the surface | [3] |
| 105 | Dr+, SSoil-, Rain- >> Phra- | Extension of the duration of the dry period is harmful for Phragmites australis | [8] |

Table S6. Model processes and constraint categories, identification number, associated formulae and descriptions of the temporary marsh ecosystem graph. The source of each rule is mentioned: [1] Common sense; [2] (Grillas et al. 2004); [3] (Grillas and Roché 1997); [4] (Lambret 2016); [5] (Chazel and Chazel 2013); [6] (Mouronval et al. 2014); [7] (Marnotte et al. 2006); [8] (Mesléard and Perennou 1996); [9] (Willm et al. 2012).

| Atmosphere | | | |
| --- | --- | --- | --- |
| 1 | TSun+, Phra-, Sci-, Junc- >> BSun+ | The opening of the vegetation allows the light to reach the bank | [3] |
| 2 | TSun+, BSun+, Zan- >> SSun+ | The opening of the vegetation allows the light to reach the surface water | [3] |
| Fauna | | | |
| 3 | Chara-, Zan- >> PeloC- | Pelobate larvae feed on algae | [2] |
| 4 | Sum+, SW- >> PeloC- | A decrease in the water level in summer leads to the death of pelobate larvae | [2] |
| 5 | Win+, SW- >> PeloC- | A decrease in the water level in winter leads to the death of pelobate larvae | [2] |
| 6 | Sum+, Sasoil+ >> PeloC- | Pelobate eggs do not support high salinity | [2] |
| 7 | SW+, Fall+ >> Lest- | Lestes macrostigma, uncompetitive, is eliminated by the other species when there is water in autumn | [4] |
| 8 | Sum+, SW+ >> Lest- | Lestes macrostigm needs the pool to dry out in summer | [4] |
| 9 | Fall+, SW+ >> Lest- | Lestes macrostigm needs the pool to dry out in fall | [4] |
| 10 | Phra-, Sci-, Junc- >> Ish- | Without refuge to hide, Ishnura pulmio is eminated by the other species of dragonflies | [2] |
| 11 | Bov+, IBG+ >> Lest-, Chara- | Bovines cause suspension of sediments that are harmful to Lestes macrostigma and the Characeae | [2, 4] |
| 12 | Eq+, IEG+ >> Lest-, Chara- | Equies cause suspension of sediments that are harmful to Lestes macrostigma and the Characeae | [2, 4] |
| 13 | SOPhra-, SOSci-, SOJunc-, Fod- >> Sus- | If there are no more resources, Sus scrofa disappears | [10] |
| 14 | Win+, SSci-, SOSci-, Chara- >> Anat- | If there are no more resources, Anatidae disappear | [10] |
| 15 | Sum+ >> Anat- | Anatidae winter in Camargue but leave in summer | [5] |
| Flora | | | |
| 16 | Bov+, IBG+ >> Chara- | Characeae are plants of low nutrient environment (bovine excreta enriche soil) | [2] |
| 17 | Eq+, IEG+ >> Chara- | Characeae are plants of low nutrient environment (equine excreta enriche soil) | [2] |
| 18 | SSun- >> Chara- | If there is insufficient light, the algae can not develop | [2] |
| 19 | BSun- >> Zan- | If there is insufficient light, the hydrophytes can not develop | [2] |
| 20 | Phra-, Sci+, Junc+ >> Junc- | Scirpus maritimus eliminates Juncus sp. | [9] |
| 21 | Phra+, Sci+, Junc+ >> Sci-, Junc- | Phragmites australis eliminates Scirpus maritimus and Juncus sp. | [9] |
| 22 | Phra+ >> SOPhra+ | Phragmites australis storage organ | [8] |
| 23 | Sci+ >> SOSci+ | Scirpus maritimus storage organ | [8] |
| 24 | Junc+ >> SOJunc+ | Juncus sp. storage organ | [8] |
| 25 | Win+ >> Dam- | Annual plant | [2] |
| 26 | TSun- >> Riel- | Riella helicophylla needs sunlight to develop | [2] |
| 27 | Brsoil+ >> Chara-, Phra-, Zan- | Charaseae, Phragmites australis and Zannichellia sp. are not tolerant to salinity | [2] |
| 28 | Sasoil+ >> Junc-, Sci-, Riel- | Juncus sp. is not tolerant to high salinity | [2, 9] |
| 29 | Sum+, SSoil-, Rain-, Wind+ >> Sci-, Dr+ | Scirpus maritimus is not tolerant to long drought | [8] |
| Human | | | |
| 30 | Phra-, Sci-, Junc-, Dam-, Riel-, Fod- >> Bov- | If there are no more resources, bovines disappears | [10] |
| 31 | Phra-, Sci-, Junc-, Fod- >> Eq- | If there are no more resources, equines disappears | [10] |
| 32 | Phra-, Sci-, Junc-, Dam-, Riel-, Fod- >> Cap- | If there are no more resources, caprines disappears | [10] |
| 33 | Phra-, Sci-, Junc-, Dam-, Riel-, Fod- >> Ov- | If there are no more resources, ovines disappears | [10] |
| 34 | SW+ >> Ov-, Cap- | Ovines and caprines cannot grazed the underwater pool | [3] |
| 35 | IBG+, Spring+ >> Bov+, Fod+ | Bovine grazing in every season for intensive bovine grazing | [10] |
| 36 | IBG+, Sum+ >> Bov+, Fod+ | Bovine grazing in every season for intensive bovine grazing | [10] |
| 37 | IBG+, Fall+ >> Bov+, Fod+ | Bovine grazing in every season for intensive bovine grazing | [10] |
| 38 | IBG+, Win+ >> Bov+, Fod+ | Bovine grazing in every season for intensive bovine grazing | [10] |
| 39 | IEG+, Spring+ >> Eq+, Fod+ | Equine grazing in every season for intensive equine grazing | [10] |
| 40 | IEG+, Sum+ >> Eq+, Fod+ | Equine grazing in every season for intensive equine grazing | [10] |
| 41 | IEG+, Fall+ >> Eq+, Fod+ | Equine grazing in every season for intensive equine grazing | [10] |
| 42 | IEG+, Win+ >> Eq+, Fod+ | Equine grazing in every season for intensive equine grazing | [10] |
| 43 | SBG+, Spring+ >> Bov+ | Bovine grazing in summer and spring for seasonal bovine grazing | [10] |
| 44 | SBG+, Sum+ >> Bov+ | Bovine grazing in summer and spring for seasonal bovine grazing | [10] |
| 45 | SBG+, Fall+ >> Bov- | Bovine grazing in summer and spring for seasonal bovine grazing | [10] |
| 46 | SBG+, Win+ >> Bov- | Bovine grazing in summer and spring for seasonal bovine grazing | [10] |
| 47 | SBEG+, Spring+ >> Bov+, Eq+ | Bovine and equine grazing in summer and spring for seasonal bovine grazing | [10] |
| 48 | SBEG+, Sum+ >> Bov+, Eq+ | Bovine and equine grazing in summer and spring for seasonal bovine grazing | [10] |
| 49 | SBEG+, Fall+ >> Bov-, Eq- | Bovine and equine grazing in summer and spring for seasonal bovine grazing | [10] |
| 50 | SBEG+, Win+ >> Bov-, Eq- | Bovine and equine grazing in summer and spring for seasonal bovine grazing | [10] |
| 51 | SMG+, Spring+ >> Bov+, Eq+, Ov+ | Bovine, equine and ovine grazing in summer and spring for seasonal bovine grazing | [10] |
| 52 | SMG+, Sum+ >> Bov+, Eq+, Ov+ | Bovine, equine and ovine grazing in summer and spring for seasonal bovine grazing | [10] |
| 53 | SMG+, Fall+ >> Bov-, Eq-, Ov- | Bovine, equine and ovine grazing in summer and spring for seasonal bovine grazing | [10] |
| 54 | SMG+, Win+ >> Bov-, Eq-, Ov- | Bovine, equine and ovine grazing in summer and spring for seasonal bovine grazing | [10] |
| Water | | | |
| 55 | Phr- >> SSoil-, SW- | The fresh phreatic water must first be filled before having water in the soil and surface water | [10] |
| 56 | SSoil- >> SW- | The soil must first be filled before having surface water | [10] |
| 57 | Rain+ >> Dr- | Rain means end of drought | [10] |

### References

Chazel, L. and M. Chazel. 2013. Camargue: Un écosystème entre terre et eau. Editions Quae.

Cohez, D., L. Paix, L. Gabrie, and A. Olivier. 2016. Plan de gestion 2016-2020 - Vol. I: Diagnostic du site (Plan de gestion). Réserve Naturelle Régionale Tour du Valat, Arles (France).

Gaucherel, C. and F. Pommereau. 2019. Using discrete systems to exhaustively characterize the dynamics of an integrated ecosystem. Methods in Ecology and Evolution **In press**.

Grillas, P., P. Gauthier, N. Yavercovski, and C. Perennou. 2004. Mediterranean Temporary Pools. Station Biologique de la Tour du Valat, Arles (France).

Grillas, P. and J. Roché. 1997. Vegetation of temporary marshes. Ecology and management. Station Biologique de la Tour du Valat. ed, Arles (France).

Lambret, P. 2016. Etude de l’écologie de Lestes macrostigma et restauration de son habitat. Courr. Nat.

Marnotte, P., A. Carrara, F. Girardot, and E. Dominati. 2006. Plantes des rizières de Camargue. Quae. ed.

Mesléard, F. and C. Perennou. 1996. Aquatic emergent vegetation. Ecology and management. Station Biologique de la Tour du Valat. ed, Arles (France).

Mouronval, J.-B., A.-L. Brochet, P. Aubry, and M. Guillemain. 2014. Les anatidés hivernant en Camargue se nourrissent-ils dans les marais aménagés pour la chasse ? Faune Sauvage **8**.

Proulx, S. R., D. E. L. Promislow, and P. C. Phillips. 2005. Network thinking in ecology and evolution. TRENDS in Ecology and Evolution **20**:345-353.

Reisig, W. 2013. Understanding Petri Nets. Springer Berlin Heidelberg, Berlin, Heidelberg.

Willm, L., N. Yavercovski, L. Mishler, and E. Mesléard. 2012. Refus de pâturage dans les parcours de Camargue. Station Biologique de la Tour du Valat.
